# PDS5A and TOP2B cooperate for chromatin recruitment via CTCF

**DOI:** 10.64898/2026.04.02.715958

**Authors:** Edgar Gonzalez-Buendia, Havva Ortabozkoyun, Danny Reinberg

## Abstract

PDS5A, a regulatory subunit of the cohesin complex, and topoisomerase IIB (TOP2B), an enzyme resolving DNA topological problems, interact with CTCF and regulate transcription, chromatin loops, and genome organization. Yet, how PDS5A and TOP2B are recruited to chromatin to exert their function is not well-understood. Here, we studied the functional relationship between PDS5A and TOP2B and the resultant impact on genome organization and gene expression. Interestingly, TOP2B-PDS5A cooperate for their recruitment to CTCF-bound chromatin sites. The presence of catalytically active TOP2B increased PDS5A occupancy genome-wide. Notably, a novel PDS5A-CTCF interaction region in the CTCF N-terminal 95-116aa was required for CTCF-PDS5A-TOP2B interaction *in vitro* as well as for active TOP2B-mediated enrichment of PDS5A chromatin occupancy *in vivo*. The loss of CTCF(95-116aa) led to a reduced number of chromatin loops and dysregulated gene expression. In gliomas, PDS5A and TOP2B expression levels are variable and correlated, contributing to apparent heterogeneity in gene expression. Indeed, inducible knockdown of PDS5A led to reduced TOP2B occupancy and altered gene expression in the glioma genome. Importantly, PDS5A mediated sensitivity to TOP2 cancer drugs in glioma cells. This newly recognized functional interaction between PDS5A and TOP2B at chromatin boundaries clarifies the mechanisms fostering gene regulation through genome organization, with implications for glioma therapeutics.

## Introduction

The organization of the mammalian genome into chromatin loops, topologically associating domains (TADs), and genome compartments contributes to appropriate gene expression profiles^1–4^. The mechanisms fostering gene regulation through three-dimensional genome organization rely on the formation and maintenance of chromatin loops, which enable the relevant interactions of DNA regulatory elements, such as enhancers and promoters^5–8^. The chromatin anchors of these loops comprise insulation factors that demarcate the discrete chromatin boundaries. Chromatin loops are formed through the action of the cohesin complex, an ATP-motor capable of establishing DNA interactions and extruding chromatin loops *in vitro* until its inevitable stalling at sites comprising either CTCF^9–12^ and/or other transcription factors with insulating capacity, such as MAZ^13,14^, PATZ1^15^, and other Zinc Finger proteins (ZNFs)^15^. Within this context, the cohesin regulators, PDS5A/B, are not required for loop extrusion *in vitro*, but instead for the resultant length of the chromatin loops *in vivo*^16–20^. Thus, PDS5A-bound cohesin maintains chromatin loops at CTCF binding sites. Yet, the recruitment of PDS5A at CTCF binding sites is not well-understood.

Changes in the chromatin loops can arise from alterations in the insulation factors, including CTCF^21–23^, MAZ^13,14^, PATZ1^15^, and other ZNFs^15^, and also in the subunits of the cohesin complex^24,25^, including PDS5A/B^16,18^. Such perturbations can impact the integrity of the chromatin domains. Thus, the processes involving these central proteins impact normal genome organization and the spatial-temporal regulation of gene expression during development, such that their disruption can foster diseases, including cancer^26–29^. Despite the advances in the study of genome organization and its impact on gene expression, our understanding of the basic mechanisms of chromatin loop formation and its maintenance remains limited.

Mounting evidence supports the loop extrusion model as a viable mechanism that continuously produces chromatin loops in an ATP-dependent fashion by loading cohesin onto chromatin fibers, thereby folding the fibers into loops^11,30,31^. Notably, cohesin-mediated loop extrusion generates DNA supercoils while the loop is protruding from the cohesin complex *in vitro*^32^. In addition, previous work has associated the supercoiling of the DNA template during transcription with genome organization^33–35^. Supercoiling is accompanied by DNA torsional stress, which is relieved by the actions of topoisomerases II A/B (TOP2A/B) that generate a transient double-strand break (DSB) and re-join both DNA strands, leading to the interconversion of topological DNA forms^36^. While TOP2A is essential for cell division, TOP2B is involved in transcription control^37,38^. Of note, TOP2B co-localizes with CTCF and cohesin at the chromatin loop anchors^39^ and its enzymatic activity can generate chromosomal rearrangements frequently found in cancers^40–44^. While the mechanisms of TOP2A/B enzymatic function have been extensively studied, their recruitment to chromatin is not well-understood.

Interestingly, TOP2A/B are overexpressed in many tumors including glioblastomas (GBM)^45^, and many available cancer drugs target TOP2A/B enzymatic function^46,47^. However, these therapies generally give rise to unpredictable and variable responses. GBM, for example, are the most malignant amongst gliomas, do not respond predictably to therapies, and exhibit heterogeneity in gene expression as well as lethality. Furthermore, TOP2B has been associated with secondary malignancies that result from TOP2 drug-induced translocations^41,48–50^. Thus, understanding the mechanisms underlying the recruitment of TOP2B activity to chromatin will very likely inform the appropriate and efficacious use of TOP2-drugs.

Here, we investigated the relationship between CTCF, PDS5A, and TOP2B. By trapping the active form of TOP2 on chromatin using a drug called Etoposide (ETO), we observed an enrichment of PDS5A at CTCF binding sites in mouse embryonic stem cells (mESCs), cervical motor neurons (MNs), and glioma cells pointing to a functional link between TOP2B activity and PDS5A localization at chromatin borders. Remarkably, the distribution of PDS5A on chromatin coincided with the binding motifs of CTCF and other insulation factors, such as MAZ, PATZ1, ZNFs, and VEZF1. We mapped a novel region of 95-116 amino-acids within the N-terminal domain of CTCF as being critical for CTCF interaction with PDS5A and TOP2B, in addition to the previously reported PDS5A-interaction sites on CTCF^17^. In gliomas, we found that PDS5A and TOP2B expression levels correlate across specimens and that TOP2B enzymatic activity is linked to PDS5A at chromatin boundaries in glioma cells. Strikingly, PDS5A knockdown reduced TOP2B occupancy in glioma cells, indicating that PDS5A occupancy might mediate sensitivity to TOP2 drugs in these cells. Our results support a model in which PDS5A is incorporated into chromatin through the enzymatic activity of TOP2B, and that PDS5A-TOP2B interaction is required for proper TOP2B occupancy genome-wide. These findings clarify the heterogenous GBM response to TOP2 drugs as well as inform their use in patients.

## Results

### CTCF, TOP2B, and PDS5A interact in mouse and human cells

Based on previous work, CTCF interacts with TOP2B and the cohesin complex at topological domain borders in HeLa cells^39^. Nonetheless, a mechanistic understanding of TOP2B recruitment to chromatin is lacking. Given that TOP2B localizes at topological borders, we investigated the connection between CTCF, TOP2B, the cohesin complex as well as its accessory subunits on chromatin. We initially performed immunoprecipitation (IP) followed by western blot (WB) analysis in HeLa nuclear extracts. Our findings indicated that CTCF co-immunoprecipitated with TOP2B and PDS5A/B in HeLa cells (Fig. 1A). We also observed RAD21 and WAPL, consistent with previous studies showing CTCF-RAD21^51^ and PDS5-WAPL interactions^52^ (Fig. 1A). In contrast to TOP2B that clearly interacts with CTCF, TOP2A does not appear to interact with CTCF in HeLa cells (Fig. 1A). Similarly, IP assays performed in nuclear extracts from mESCs indicated that CTCF interacted specifically with TOP2B, compared to TOP2A (Fig. 1B). Interestingly, both TOP2A and TOP2B interact with each other yet the expression levels of TOP2A are higher than those of TOP2B in mESCs (see input, Fig. 1B). Taken together, TOP2B interacts with PDS5A/B and CTCF.

**Fig. 1.**
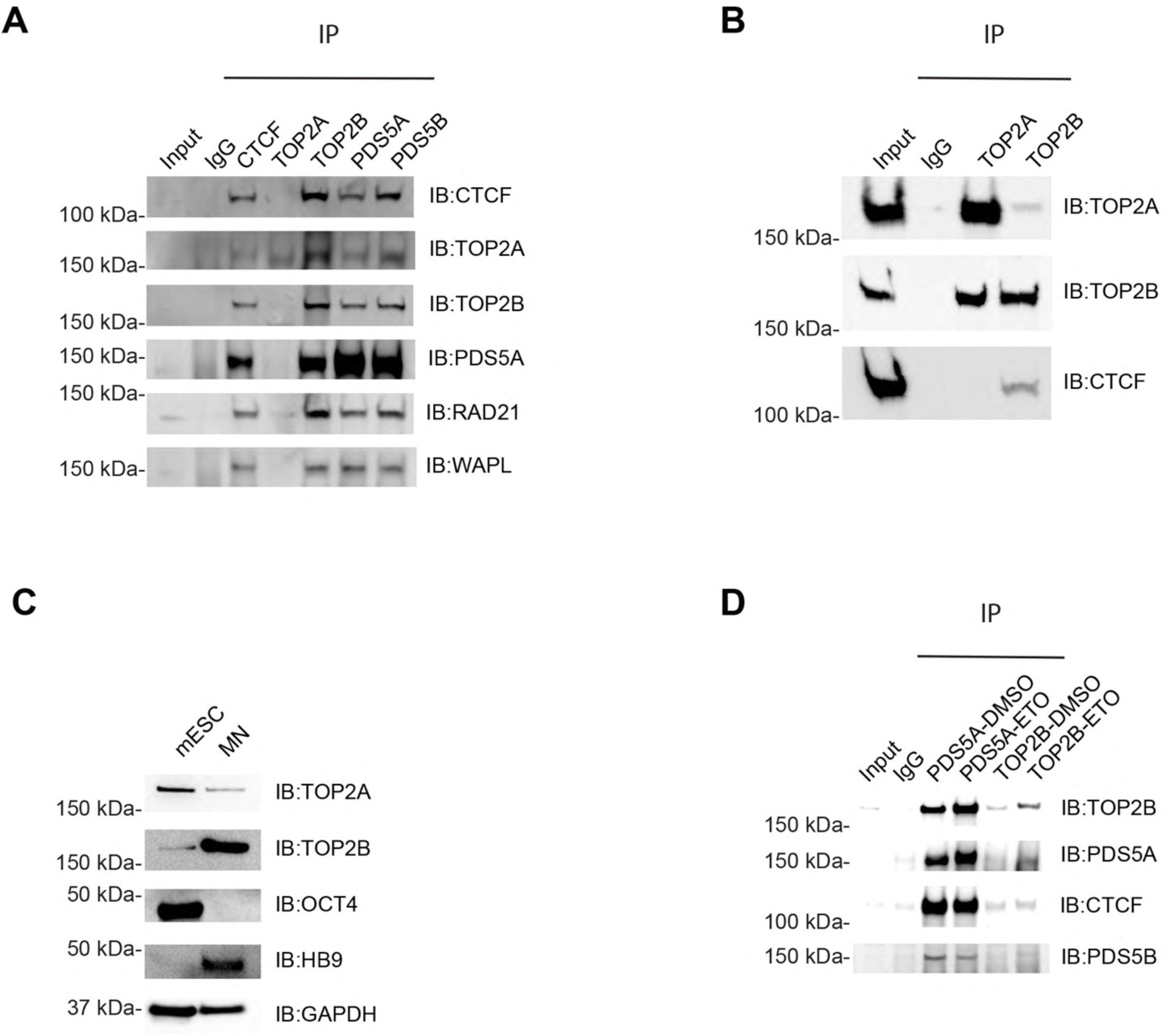
TOP2B and PDS5A interact with CTCF in human and mouse cells. **(A)** CTCF, TOP2B, and PDS5A/B interaction in human HeLa cells, as determined by Immunoprecipitation (IP) of CTCF, TOP2A, TOP2B, PDS5A, and PDS5B; one of two independent IP experiments using chromatin fractions after conventional chromatography fractionation on a phosphocellulose column. Western blot analysis (WB) of CTCF, TOP2A, TOP2B, PDS5A, RAD21, and WAPL were performed for each IP. IgG serves as a control. IB: Immunoblot. **(B)** TOP2B interacts with CTCF in mESC as determined by IP and WB; two independent IP experiments of TOP2A and TOP2B using the Dignam protocol for preparation of nuclear extract protein. TOP2A IP did not reveal any interaction with CTCF. IgG serves as a control. IB: Immunoblot. **(C)** TOP2B protein levels are higher in motor neurons (MNs) differentiated from mESCs than in mESCs as determined by WB. TOP2A protein levels decrease upon differentiation. OCT4 serves as a marker for mESCs, HB9 for MNs, and GAPDH serves as a loading control; two independent experiments using protein preparations from whole cell extracts. IB: Immunoblot. **(D)** PDS5A exhibits increased chromatin binding upon Etoposide (ETO) trapping of TOP2B relative to the control (DMSO) in MNs. PDS5A and TOP2B levels in MNs derived from mESCs were gauged by IP and WB. CTCF interaction with either PDS5A or TOP2B on chromatin was similar irrespective of ETO treatment, as evidenced by WB. PDS5A IP upon ETO treatment did not exhibit increased interaction with PDS5B on chromatin. TOP2B IP did not reveal any interaction with PDS5B. Cells were treated with 0.5 mM ETO for 10 min for TOP2 trapping on chromatin. IB: Immunoblot.

Given that PDS5A/B regulate chromatin loops at CTCF binding sites^16,18^, we sought to determine whether these proteins work together to recruit TOP2B to chromatin. As we observed that the pharmacological induction of TOP2B enzymatic activity influences TOP2B localization in the genome across human glioma cell lines^45^, we investigated TOP2B interaction with PDS5A/B using Etoposide (ETO), a TOP2 poison that traps “active” TOP2A/B on chromatin. This process of TOP2 trapping occurs after the TOP2-elicited DNA double-strand break (DSB) and before its DNA re-joining^46^. As TOP2A and CTCF interaction was not detectable in mESCs (Fig. 1B), we aimed to discriminate TOP2A from TOP2B enzymatic activity using differentiated motor neurons (MNs) derived from mESCs, wherein TOP2A expression levels range from low to undetectable as a result of differentiation towards post-mitotic neurons (Fig. 1C). As expected, based on the switch from TOP2A to TOP2B isoforms during neuronal differentiation^53^, higher levels of TOP2B were observed in MNs compared to those in mESCs (Fig. 1C). We next isolated the chromatin from MNs, either untreated or treated with ETO to trap “active” TOP2B on chromatin, and performed IP assays for PDS5A and TOP2B. PDS5A was enriched on chromatin from ETO-treated MNs as compared to non-treated cells (ETO vs DMSO, respectively, Fig. 1D), as evidenced by WB analysis of TOP2B and PDS5A IPs. In contrast, ETO-treatment was ineffectual with respect to the levels of CTCF, PDS5B, and other cohesin-associated components, such as WAPL and RAD21 (Fig. 1D and see fig. S1A for WAPL and RAD21). This data points to a previously unrecognized functional link between TOP2B and PDS5A: the trapped, active TOP2B and/or the product of its activity (*i.e.* DSB) promotes an increase of PDS5A on chromatin. Notably, we had previously detected PDS5A within the top 20 candidates whose loss further impacts the integrity of the *Hoxa5|6* boundary in MNs already having a deletion of a CTCF DNA-binding site (CTCF Δ5|6:6|7)^14^. Taken together, TOP2B and PDS5A interact with CTCF on chromatin, and PDS5A levels were increased upon trapping active TOP2B. These results prompted us to investigate PDS5A localization as a function of ETO treatment using ChIP-seq.

### TOP2B enzymatic activity enriches PDS5A at chromatin boundaries dependent on the presence of CTCF

Previous work in mESCs showed that PDS5A specifically interacts with CTCF^17^, and cooperates with PDS5B in regulating chromatin loops^16,18^. Thus, we next probed for the relationship between CTCF and PDS5A/B upon ETO treatment to “trap” TOP2A/B on chromatin using ChIP-seq in mESCs. We used mESCs having a CTCF-degron (CTCF-GFP-AID-Tir1)^22^, either untreated (CTCF-WT) or treated with auxin to degrade endogenous CTCF (ΔCTCF), followed by DMSO versus ETO treatment before ChIP-seq. Importantly, ETO treatment did not affect CTCF protein levels (fig. S1B) or CTCF occupancy on chromatin (Fig. 2A-C). Moreover, we and others have observed previously that the cohesin subunit, RAD21, re-localizes on chromatin in the absence of CTCF in mESCs^15^ and this outcome also occurred irrespectively of ETO treatment (Fig. 2D-F), suggesting that TOP2 activity does not affect cohesin complex localization. Similar to our previous IP results (Fig. 1D), ETO treatment was ineffectual with respect to PDS5B binding on chromatin by ChIP-seq (Fig. 2G-I). However, the absence of CTCF did lead to the loss of PDS5B, compared to the CTCF-WT case (Fig. 2G-I). In contrast to PDS5B, PDS5A remained bound on chromatin upon CTCF degradation (ΔCTCF, Fig. 2J-L and fig. S1C). Remarkably, ETO-mediated chromatin trapping of TOP2B led to enriched PDS5A occupancy genome-wide (Fig. 2J-L), as also observed in TOP2B or PDS5A IPs from chromatin (Fig. 1D). Interestingly, this enrichment was connected to the presence of CTCF (CTCF-WT) as evidenced by its abolishment upon CTCF degradation (ΔCTCF, Fig. 2J-L).

**Fig. 2.**
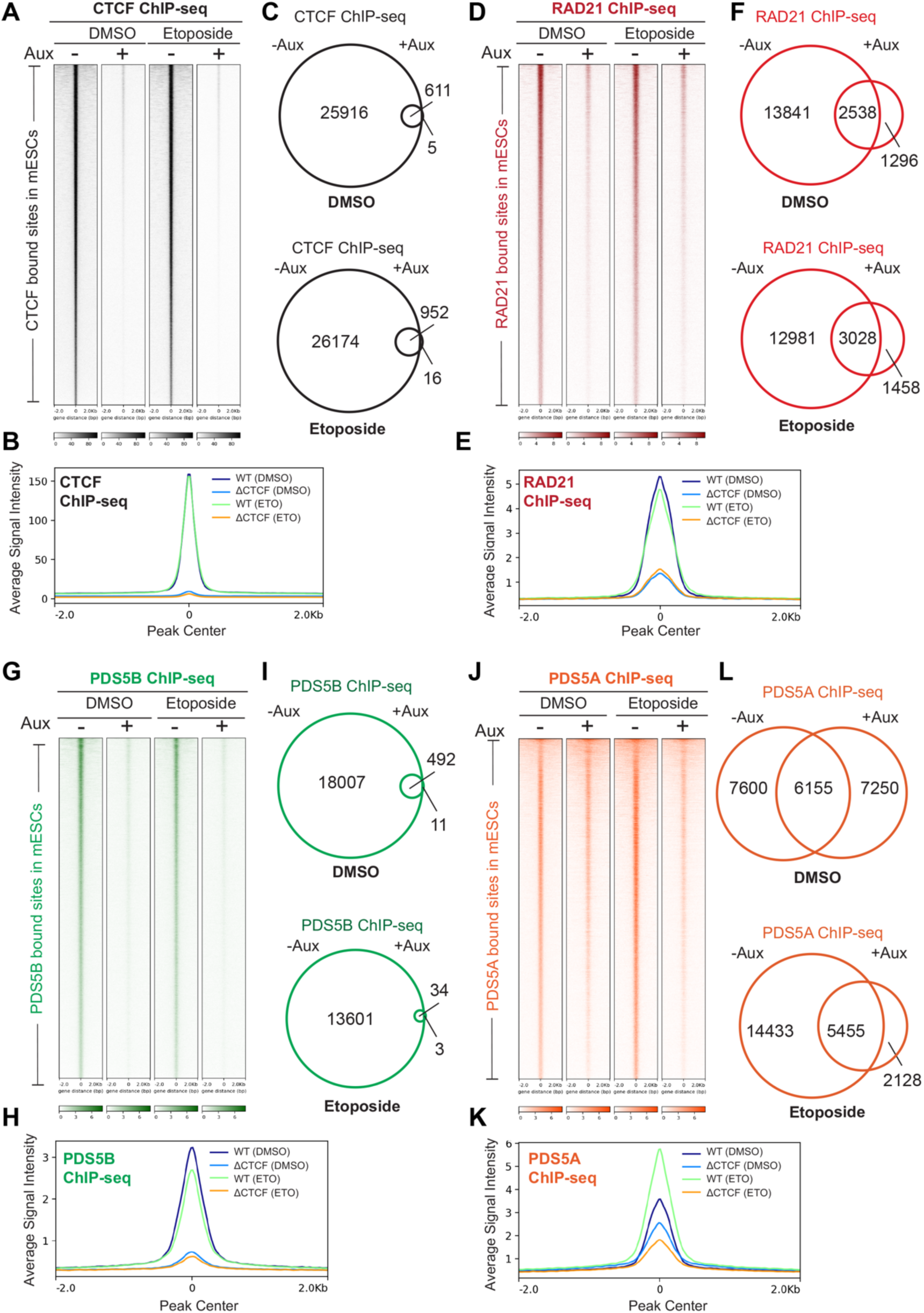
PDS5A binding is enhanced by TOP2 activity on chromatin in the presence of CTCF in mESCs. **(A)** Heat maps showing CTCF ChIP-seq read densities in mESCs under the indicated conditions in CTCF-degron mESCs: DMSO or ETO treatment conditions in CTCF intact (WT, -Aux) or CTCF degraded (ΔCTCF, +Aux) mESCs. Cells were treated with auxin for 48 hr to degrade CTCF. Cells were treated with 0.5 mM ETO for 10 min for TOP2 trapping on chromatin. ChIP-seq data represents one representative biological replicate for each condition. **(B)** Average density profile plots of CTCF ChIP-seq under conditions of DMSO versus ETO treatment in WT or ΔCTCF mESCs. **(C)** Venn diagrams indicating CTCF binding in WT versus ΔCTCF mESCs under DMSO or ETO conditions. **(D)** Heat maps showing RAD21 ChIP-seq read densities in mESCs under the indicated conditions in CTCF-degron mESCs. ChIP-seq data represents one representative biological replicate for each condition. **(E)** Average density profile plots of RAD21 ChIP-seq under the conditions of DMSO versus ETO treatment in WT or ΔCTCF mESCs. **(F)** Venn diagrams indicating RAD21 binding in WT versus ΔCTCF mESCs under DMSO or ETO conditions. **(G)** Heat maps showing PDS5B ChIP-seq read densities in mESCs under the indicated conditions in CTCF-degron mESCs. ChIP-seq data is from one representative of two biological replicates for each condition. **(H)** Average density profile plots of PDS5B ChIP-seq under the conditions of DMSO versus ETO treatment in WT or ΔCTCF mESCs. **(I)** Venn diagrams indicating PDS5B binding in WT versus ΔCTCF mESCs under DMSO or ETO conditions. **(J)** Heat maps showing PDS5A ChIP-seq read densities in mESCs under the indicated conditions in CTCF-degron mESCs. ChIP-seq data is from one representative of two biological replicates for each condition. **(K)** Average density profile plots of PDS5A ChIP-seq under the conditions of DMSO versus ETO treatment in WT or ΔCTCF mESCs. **(L)** Venn diagrams indicating PDS5A binding in WT versus ΔCTCF mESCs under DMSO or ETO conditions.

While PDS5A was localized mostly at promoters, intronic and intergenic regions in CTCF-WT cells, its localization at promoters was increased in ΔCTCF cells (fig. S1D). In addition to this global analysis, we also observed examples of individual loci bound by PDS5A in the absence of CTCF. For example, PDS5A localizes with CTCF in the *HoxA* cluster and remains bound on chromatin even in the absence of CTCF (fig. S2). Notably, the binding pattern of PDS5A at the *HoxA* cluster is consistent with our previous observation that its loss further disrupts the *Hoxa5|6* boundary in MNs lacking CTCF binding at the *Hoxa5|6* boundary^14^.

To further analyze this enhanced PDS5A chromatin binding upon ETO treatment, we compared PDS5A ChIP-seq peaks under conditions of DMSO versus ETO treatment (fig. S3). TOP2B activity induced by ETO appears to incorporate PDS5A at 7438 *de novo* binding regions in addition to enriching the vast majority of the 12512 PDS5A peaks observed under DMSO conditions (fig. S3A). To gain insight into these PDS5A-enriched genomic regions, we ascertained the DNA-motifs to which PDS5A localizes upon ETO treatment and assessed the nuclear factors that bind to these motifs (fig. S3B). Remarkably, CTCF as well as other insulation factors such as MAZ, PATZ1, ZNFs, and VEZF1 matched to the motifs overrepresented at all of these ETO-enriched sites, whether novel or present with DMSO, thereby linking TOP2B activity and PDS5A localization to chromatin insulators. Taken together, these results show that PDS5A enrichment at chromatin boundaries is not only controlled by the presence of active TOP2B or the product of its activity (*i.e.* DSB) in a CTCF-dependent manner, but this enrichment is localized to motifs associated with CTCF and other insulation factors.

### A novel PDS5A-CTCF interaction region facilitates PDS5A-TOP2B localization at chromatin boundaries

Given the requirement of CTCF for PDS5A enrichment, we investigated CTCF-PDS5A interaction using alpha fold protein-protein prediction tools to ascertain the CTCF-PDS5A interaction surfaces. In addition to the previously reported PDS5A-interacting region on the CTCF N-terminal 13-33 amino acids (region 1)^17^, we found a novel CTCF-PDS5A interacting region on the CTCF N-terminal amino acids 95-116 (region 2; fig. S4A-B). To understand the impact of these two CTCF regions on PDS5A interaction on chromatin, we performed rescue experiments in auxin-treated ΔCTCF background mESCs using HA-FLAG-tagged versions of CTCF, either WT or having a deletion in either region 1 [MUT1(Δ13-33)] or region 2 [MUT2(Δ95-116)]. (Fig. 3A). Endogenous CTCF-GFP was degraded under these conditions, as expected, and the different CTCF versions rescued CTCF to similar levels as assessed by WB of nuclear extracts (Fig. 3B). Of note, TOP2B and PDS5A expression seemed to be regulated by CTCF as auxin treatment reduced their levels, as evident in the case of RAD21 as well (Fig. 3B). Importantly, the expression of PDS5A, TOP2B, and RAD21 were restored to levels similar to those of control cells upon expressing any of the HA-FLAG-CTCF constructs: CTCF-WT or CTCF-MUT1(Δ13-33) or CTCF-MUT2(Δ95-116) (Fig. 3B).

**Fig. 3.**
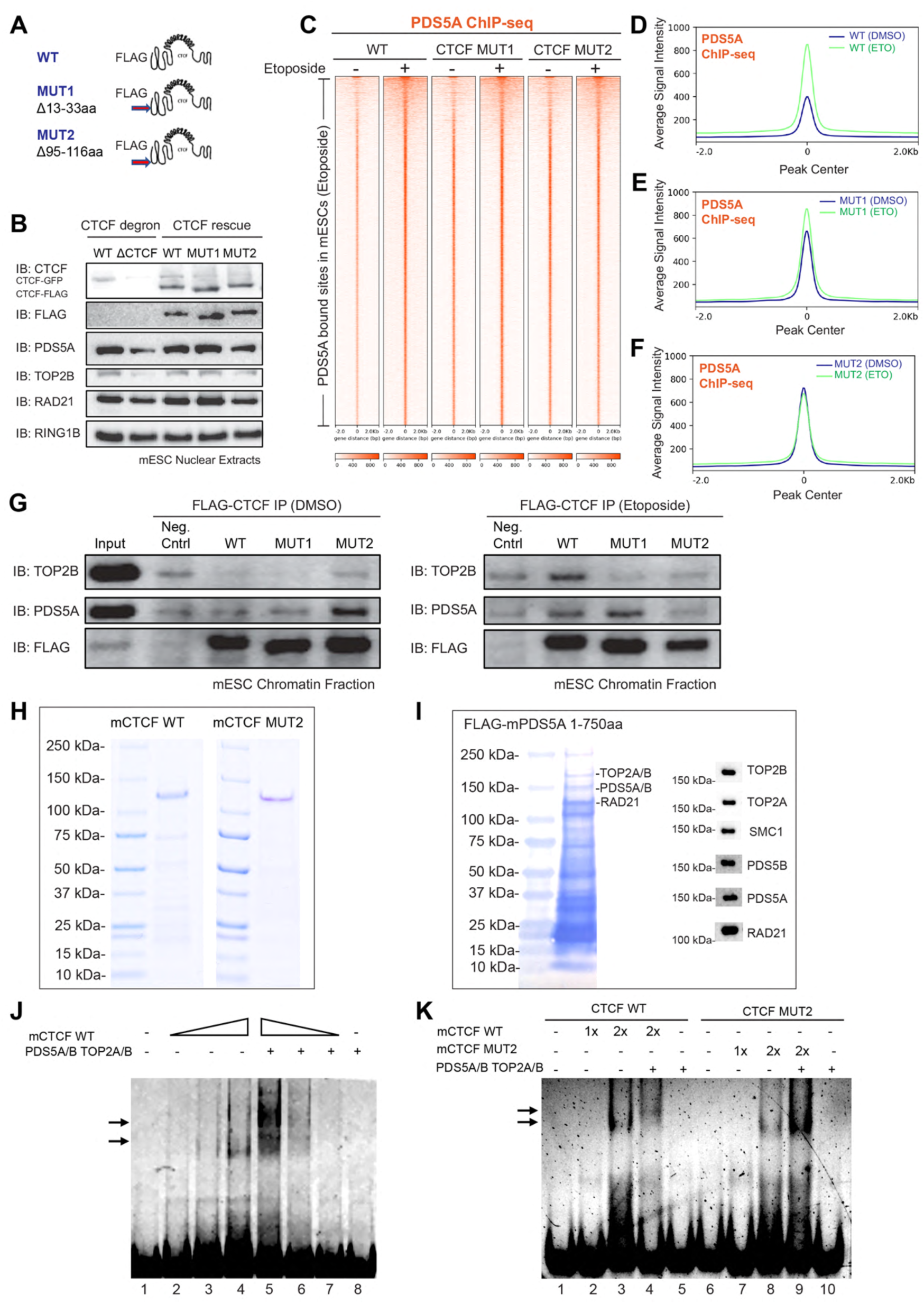
A novel PDS5A-CTCF interaction region in the CTCF N-terminus (95-116aa) is required for PDS5A enrichment at CTCF bound sites. **(A)** Schematic showing WT and mutant versions of CTCF, as indicated. **(B)** Different versions of HA-FLAG-CTCF N-terminus mutants were ectopically expressed in CTCF-degron mESCs treated with auxin to eliminate endogenous GFP-tagged CTCF (ΔCTCF). *Topmost panel*: WB using CTCF antibody to detect HA-FLAG-CTCF comprising either full-length (FL) CTCF (WT), CTCF MUT1(Δ13-33), or CTCF MUT2(Δ95-116) expressed in ΔCTCF mESC background (CTCF-GFP untreated, -Aux). *Bottom panels:* Protein expression levels of versions of FLAG-tagged CTCF, PDS5A, TOP2B, RAD21, and RING1B detected by WB. RING1B served as loading control. **(C)** Heat maps showing PDS5A ChIP-seq read densities at PDS5A-bound sites in mESCs expressing HA-FLAG-CTCF-WT (WT), HA-FLAG-CTCF-MUT1(Δ13-33) (CTCF-MUT1), and HA-FLAG-CTCF-MUT2(Δ95-116) (CTCF-MUT2) under conditions of DMSO versus ETO treatment. ChIP-seq data is from one representative biological replicate for each condition. **(D-F)** Average density profile plots showing PDS5A ChIP-seq read densities under conditions of DMSO versus ETO treatment in mESCs expressing CTCF, either WT (**D**), CTCF-MUT1 (**E**), or CTCF-MUT2 (**F**). ChIP-seq data is from one representative biological replicate for each condition. **(G)** Interaction between TOP2B, PDS5A, and CTCF as a function of the presence of the CTCF N-terminus mutants. WB analysis of FLAG-IP performed on HA-FLAG-tagged versions of the CTCF mutants expressed in a ΔCTCF mESC background under the conditions of DMSO (left panel) or ETO treatment (right panel), as indicated (one of two independent FLAG-IP experiments using Chromatin fraction; Dignam Nuclear pellet preparation was treated with Benzonase nuclease). **(H)** Purified FLAG-mCTCF-WT and FLAG-mCTCF-mutant version (MUT2) recombinant mouse proteins. **(I)** Purified FLAG-mPDS5A (1-750aa) recombinant protein, which co-purified with TOP2A/B, PDS5A/B, and cohesin components as shown by Coomassie blue staining (left), and WB analysis (right). **(J)** Electrophoretic Mobility Shift Assay (EMSA) indicating *in vitro* reconstitution of CTCF-PDS5A-TOP2B interaction. Arrows indicate the shifted migration of the probe upon addition of CTCF alone and the further shifting of the CTCF-bound probe upon addition of PDS5 and TOP2B. The experiment was performed as a function of increasing or decreasing CTCF levels, as indicated. **(K)** EMSA indicating that while both WT and MUT2 versions of CTCF bind to the probe containing the CTCF motif, only CTCF-WT participates in CTCF-PDS5-TOP2 association (n=2).

To test whether the deletion mutants affect CTCF interaction with PDS5A or prevent PDS5A enrichment at CTCF boundaries via TOP2B activity, we treated the CTCF rescue cell lines with DMSO or ETO and performed ChIP-seq with HA (fig. S4C) or PDS5A (Fig. 3C-F and fig. S5A-D). In CTCF-WT cells, we observed a gain of PDS5A genome-wide both at CTCF binding sites (fig. S5A-B) and at PDS5A binding sites (Fig. 3C-D) upon ETO treatment, as expected. In the case of CTCF-MUT1(Δ13-33), a weak PDS5A enrichment upon ETO treatment was observed genome-wide, as compared to CTCF-WT (Fig. 3C-E and fig. S5A-C). In contrast, CTCF-MUT2(Δ95-116) failed to enrich for PDS5A upon ETO treatment genome-wide, as compared to CTCF-WT (Fig. 3C, F and fig. S5A, D). Importantly, CTCF-WT, CTCF-MUT1(Δ13-33) and CTCF-MUT2(Δ95-116) exhibited similar levels of DNA binding in untreated cells (DMSO) as a control (fig. S4C). While we observed a modest decrease in CTCF-WT binding upon ETO treatment, CTCF-MUT1(Δ13-33) and CTCF-MUT2(Δ95-116) showed similar levels of CTCF binding irrespective of ETO treatment (fig. S4C).

To verify the protein interactions on chromatin, we performed CTCF FLAG-IP followed by WB for FLAG-CTCF, PDS5A, and TOP2B using cells expressing CTCF-WT, CTCF-MUT1(Δ13-33) or CTCF-MUT2(Δ95-116), either untreated (DMSO) or under ETO-treatment conditions (Fig. 3G). Importantly, CTCF-WT co-immunoprecipitated with PDS5A and TOP2B under ETO-treated conditions (Fig. 3G, right panel), pointing to CTCF-PDS5A-TOP2B interaction in the cells. Interestingly, under similar conditions, CTCF-MUT1(Δ13-33) co-immunoprecipitated PDS5A, but not TOP2B (Fig. 3G, right panel), indicating that CTCF-MUT1(Δ13-33) impacts the interaction between TOP2B and CTCF on chromatin upon Etoposide treatment. On the other hand, CTCF-MUT2(Δ95-116) exhibited the highest levels of PDS5A co-immunoprecipitation in untreated cells (Fig. 3G, left panel), while under ETO conditions, CTCF-MUT2(Δ95-116) affected the interaction between CTCF, PDS5A, and TOP2B, as compared to CTCF-WT (Fig. 3G, right panel), consistent with our ChIP-seq results. Thus, while CTCF MUT2(Δ95-116) accumulated PDS5A on chromatin (DMSO), its interaction with PDS5A was impaired upon ETO treatment thereby preventing the chromatin-localized gain of PDS5A. Thus, active TOP2B enhances PDS5A occupancy genome-wide as a function of the presence of region 2 (95-116aa) of the CTCF-N terminus.

### The N-terminus of CTCF harboring a novel CTCF-PDS5A interaction region is required for association with PDS5A-TOP2B in vitro

To further confirm the CTCF-PDS5A-TOP2B interaction and the impact of CTCF-MUT2(Δ95-116) on PDS5A-TOP2B recruitment, we purified recombinant mouse mCTCF-WT and mCTCF-MUT2(Δ95-116) (Fig. 3H). In addition, we expressed a recombinant fragment of mouse PDS5A(1-750aa) including the conserved module that interacts with RAD21 and SMC3 *in vitro*^54,55^ and purified this protein from nuclei using FLAG affinity purification under conditions conducive to retaining its interacting partners upon elution (Fig. 3I). Indeed, WB analysis demonstrated the presence of full-length (FL) PDS5A, PDS5B, TOP2A, TOP2B, and RAD21 (Fig. 3I). We next tested whether the PDS5A-TOP2B fraction can interact with CTCF-WT protein bound to a probe containing a CTCF binding site^56^ in an Electrophoretic Mobility Shift Assay (EMSA) (Fig. 3J-K). Increasing amounts of CTCF-WT gave rise to a shift in the probe mobility (Fig. 3J, lanes 2-4), and the addition of the fraction containing PDS5A-TOP2B gave rise to another shift of slower-mobility (Fig. 3J, lane 5). To verify that the PDS5A-TOP2B-associated shift depends on CTCF-WT, we probed for the effect if any of using decreasing amounts of CTCF-WT and observed the gradual loss of both shifts corresponding to CTCF-WT and CTCF-WT-PDS5A-TOP2B (lanes 6-8) (Fig. 3J). Using the same EMSA conditions, we next tested whether PDS5A-TOP2B can bind to CTCF-MUT2(Δ95-116) (Fig. 3K). Again, increasing amounts of CTCF-WT gave rise to a shift (lane 2,3), and the addition of the fraction containing PDS5A-TOP2B gave rise to a further shift (lane 4) that was dependent on the presence of CTCF-WT (lane 5) (Fig. 3K). However, while increasing amounts of CTCF-MUT2(Δ95-116) gave rise to a shift of the probe, as expected (lanes 7,8), the addition of the fraction containing PDS5A-TOP2B fraction was ineffectual (lanes 6-10) (Fig. 3K), pointing to the inability of CTCF-motif-bound CTCF-MUT2(Δ95-116) to recruit PDS5A-TOP2B *in vitro*. Thus, CTCF amino acids 95-116 at the N-terminus interact with PDS5A-TOP2B and an intact CTCF N-terminus is required for PDS5A-TOP2B binding to CTCF *in vitro*.

### CTCF N-terminus Δ95-116aa mutant impacts genome organization and gene expression in mESCs

Based on our results described above, the recruitment of PDS5A and TOP2B to chromatin is dependent on an intact CTCF N-terminus region (Fig. 3), and possibly other insulation factors given the remaining chromatin-bound PDS5A in ΔCTCF (see Fig. 2J-L, and fig. S2-3) and its occupancy at insulator motifs in the presence of active TOP2B (fig. S3B). In addition, PDS5A occupancy on chromatin appears to be dynamic as we observed variable levels depending on TOP2B catalytical activity (Fig. 2-3). As the N-terminus of CTCF (amino acids 95-116) is required for PDS5A-TOP2B binding *in vitro* (Fig. 3J-K) and this requirement is also detectable in mESCs under conditions of ETO-treatment (Fig. 3G, right panel), we next examined the impact of CTCF-MUT2(Δ95-116aa) on genome organization via Micro-C (Fig. 4 and fig. S6). While notable changes in active and inactive compartments were not detected, as expected (fig. S6A), differential loop analysis indicated alterations in looping interactions (fig. S6B). Importantly, we observed that CTCF-MUT2(Δ95-116) led to reduced looping interactions relative to WT by Aggregate Peak Analysis (APA) (fig. S6C). Similarly, the looping interactions were reduced in CTCF-MUT2(Δ95-116) at loop anchors as categorized by the occupancy of CTCF, PDS5A, and CTCF-PDS5A (Fig. 4A and see fig. S6D for CTCF-PDS5A-RAD21-containing loops). Two example loci with reduced looping interactions in CTCF-MUT2(Δ95-116) as compared to WT are depicted in Fig. 4B-C. In addition to the impact on genome organization, CTCF-MUT2(Δ95-116) affected global gene expression in mESCs. Comparison of gene expression from CTCF-WT versus CTCF-MUT2(Δ95-116) by RNA-seq gave evidence of 1354 differentially expressed genes (DEGs, padj < 0.05) in the latter case (Fig. 4D). These DEGs corresponded to biological processes involved in development, cell signaling, and metabolism (Fig. 4E), and ∼40% of them were found in proximity to PDS5A ChIP-seq peaks in CTCF-MUT2 (Fig. 4F).

**Fig. 4.**
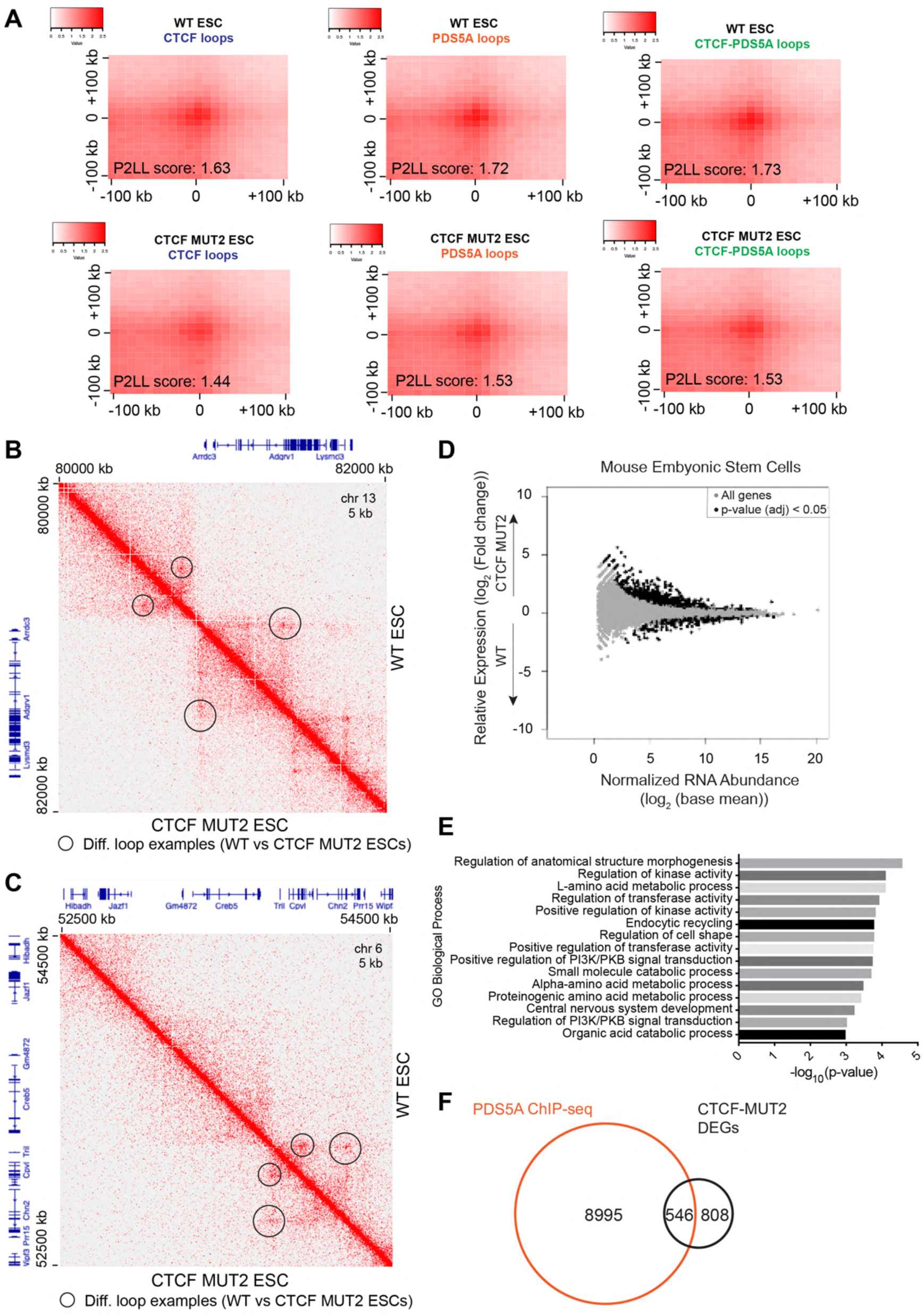
Deletion of 95-116aa of the CTCF N-terminus (CTCF-MUT2) impacts genome organization and gene expression in mESCs. **(A)** Decrease of loops in Aggregate Peak Analysis (APA) of CTCF-MUT2 compared to CTCF-WT by Micro-C assay. APA of loops were plotted in WT vs MUT2 mESCs showing ChIP-seq signals of CTCF, PDS5A, or both at any region covered by them. The resolution of APA is 10 kb. Micro-C assay represents two biological replicates in each condition. **(B-C)** Visualization of Micro-C contact matrices were depicted for example loci indicating differential loops between WT vs MUT2. **(D)** RNA-seq MA plot of CTCF-WT vs CTCF-MUT2 showing differentially expressed genes (DEGs) in mESCs from four biological replicates. **(E)** Gene Ontology analysis of DEGs in CTCF-WT vs CTCF-MUT2 RNA-seq in mESCs. **(F)** Venn diagram indicating the overlap between CTCF-MUT2 DEGs and PDS5A ChIP-seq binding in mESCs. PDS5A ChIP-seq peaks were annotated to the genes using PAVIS tools.

As we observed that CTCF-MUT1(Δ13-33) also affected TOP2B interaction in the context of CTCF-PDS5A-TOP2B (Fig. 3G, right panel), we analyzed gene expression in CTCF-WT versus CTCF-MUT1(Δ13-33) using RNA-seq (fig. S7A); 2340 genes were differentially expressed in the latter case (padj < 0.05, see fig. S7A). Interestingly, gene ontology analysis showed that mainly ribosome biogenesis pathways were affected (fig. S7B). Thus, a different set of genes were affected in CTCF-MUT1(Δ13-33) as compared to CTCF-MUT2(Δ95-116), confirming that these mutants are not redundant (fig. S7C). Taken together, our results show that CTCF-MUT2(Δ95-116), which cannot bind to PDS5A-TOP2B *in vitro,* led to reduced loop formation and altered gene expression in mESCs *in vivo,* thereby possibly impacting the expression of genes that are frequently dysregulated in cancer.

### In gliomas, PDS5A and TOP2B protein levels are correlated across tumors

Glioblastomas (GBM) are the most lethal amongst brain tumors and exhibit heterogeneity in gene expression. We previously showed that in a subset of gliomas, TOP2B is overexpressed and thereby modulates oncogene expression and tumor growth^45^. Given the functional link described herein between TOP2B and PDS5A in mESCs, and their association in gene expression and genome organization, we next tested whether TOP2B and PDS5A expression levels are correlated in gliomas. Indeed, immunohistochemistry of a brain tumor tissue array including normal samples and all tumor grade specimens in adults indicated correlative levels of TOP2B and PDS5A (Fig. 5A-C, see fig. S8 for controls), which support their interaction in gliomas and suggest a possible functional association in tumors. Moreover, we used GlioVis data portal for visualization and analysis of brain tumor expression datasets^57^. Using the CGGA dataset with 1019 RNA-seq samples from all glioma tumor grades in adults, we found that TOP2B and PDS5A expression correlated in gliomas (Pearson’s correlation r=0.72, P<0.001) (Fig. 5D-E). In addition to their correlative RNA expression levels, we used cBioportal^58–60^ to analyze a GBM dataset with 99 adult patient samples with protein quantification data determined by Mass-Spectrometry^61^. Consistent with the RNA expression levels, we found that TOP2B and PDS5A protein abundance are correlated in GBM (Pearson coefficient r=0.66, p-value=6.91e-14) (Fig. 5F). We next assessed the protein levels of TOP2B and PDS5A in the glioma cell lines via WB and found that both are higher in U87MG and BT142 cells, as compared with controls (Fig. 5G). As BT142 glioma cells expressed the highest levels of TOP2B and PDS5A and additionally, RAD21 appears to be elevated in these cells, relative to U87MG and A172 (Fig. 5G), we next investigated the potential functional association between TOP2B and PDS5A in glioma using BT142 cells.

**Fig. 5.**
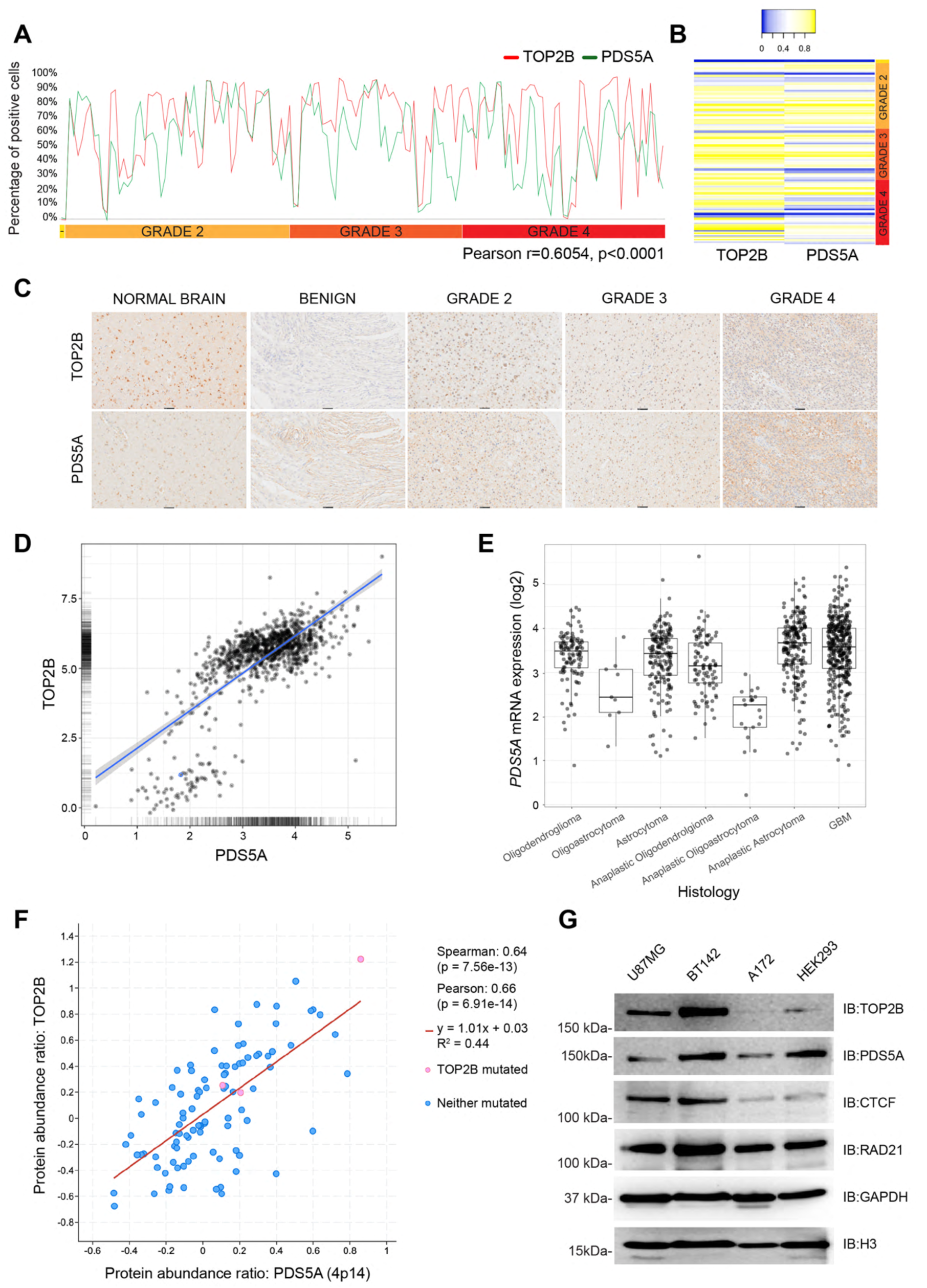
Expression of PDS5A and TOP2B is variable and correlated in human gliomas. **(A)** Correlation of TOP2B versus PDS5A expression by immunohistochemistry in human gliomas. All tumor grades in adults are shown including low-grade gliomas (grade 2 in yellow, grade 3 in orange) and GBM (grade 4 in red). Light yellow region indicates malignant meningioma. The correlation is determined by Pearson coefficient r=0.6054, p-value < 0.0001 in n=208 samples that consist of meningioma, astrocytoma and glioblastoma. Brain tumor tissue array was stained with TOP2B or PDS5A antibodies. Visiopharm software was used to detect DAB chromogen (3,3’-diaminobenzidine) and count the positive cells as percentage. **(B)** Heatmap of the percentage of positive cells in the brain tumor tissue array shown in panel (**A**), indicating the correlative expression of TOP2B and PDS5A detected by immunohistochemistry. Brain tumor tissue array was stained with TOP2B or PDS5A antibodies. Visiopharm software was used for analysis. **(C)** Immunohistochemistry of the representative brain or tumor specimens showing correlative levels of TOP2B and PDS5A in normal brain, low-grade glioma (grade 2 and 3), and GBM (grade 4). **(D)** Correlation of TOP2B versus PDS5A gene expression by RNA-seq in human gliomas. All tumor grades in adult patient samples including low-grade gliomas and GBM are from CGGA GlioVis portal. The correlation is determined by Pearson coefficient r=0.72, p-value < 0.001 in n=1019 samples. **(E)** Box plot of PDS5A expression by RNA-seq in human gliomas. All tumor grades in adults including low-grade gliomas and GBM are from GlioVis portal (n=1019 samples). **(F)** Correlation of TOP2B versus PDS5A protein abundance determined by Pearson coefficient r=0.66, p-value=6.91e-14. The datasets of adult patient samples across all tumor grades are from cBioportal. **(G)** Expression of TOP2B, PDS5A, CTCF, RAD21, GAPDH, and histone H3 in glioma cell lines (U87MG, BT142, and A172), and Human Embryonic Kidney cell line (HEK293) by WB. GAPDH and H3 served as loading controls. IB: Immunoblot.

### PDS5A recruitment is enhanced by TOP2B activity at CTCF binding sites in BT142 glioma cells

We first examined TOP2B and PDS5A co-localization at chromatin boundaries in BT142 glioma cells after DMSO or ETO-treatment followed by ChIP-seq for CTCF, PDS5A, and TOP2B (Fig. 6). As previously observed in mESCs, CTCF binding is not affected by ETO treatment (Fig. 6A, left). Remarkably, we observed a gain in PDS5A under conditions of ETO, but not DMSO treatment (Fig. 6A-D). Similar to our results shown above (Fig. 2J-L), the binding sites of PDS5A overlapped those of CTCF (Fig. 6B). Thus, the enrichment in PDS5A binding following TOP2 trapping on chromatin (ETO) was localized mainly to CTCF binding sites (Fig. 6A-D), and TOP2B co-localized with both CTCF and PDS5A across these regions (Fig. 6A-D). Interestingly, TOP2B was also observed at other locations without CTCF (Fig. 6E and fig. S9). In other words, CTCF, PDS5A, and TOP2B co-localization was evident in ∼27 % of TOP2B-bound sites (see Fig. 6B).

**Fig. 6.**
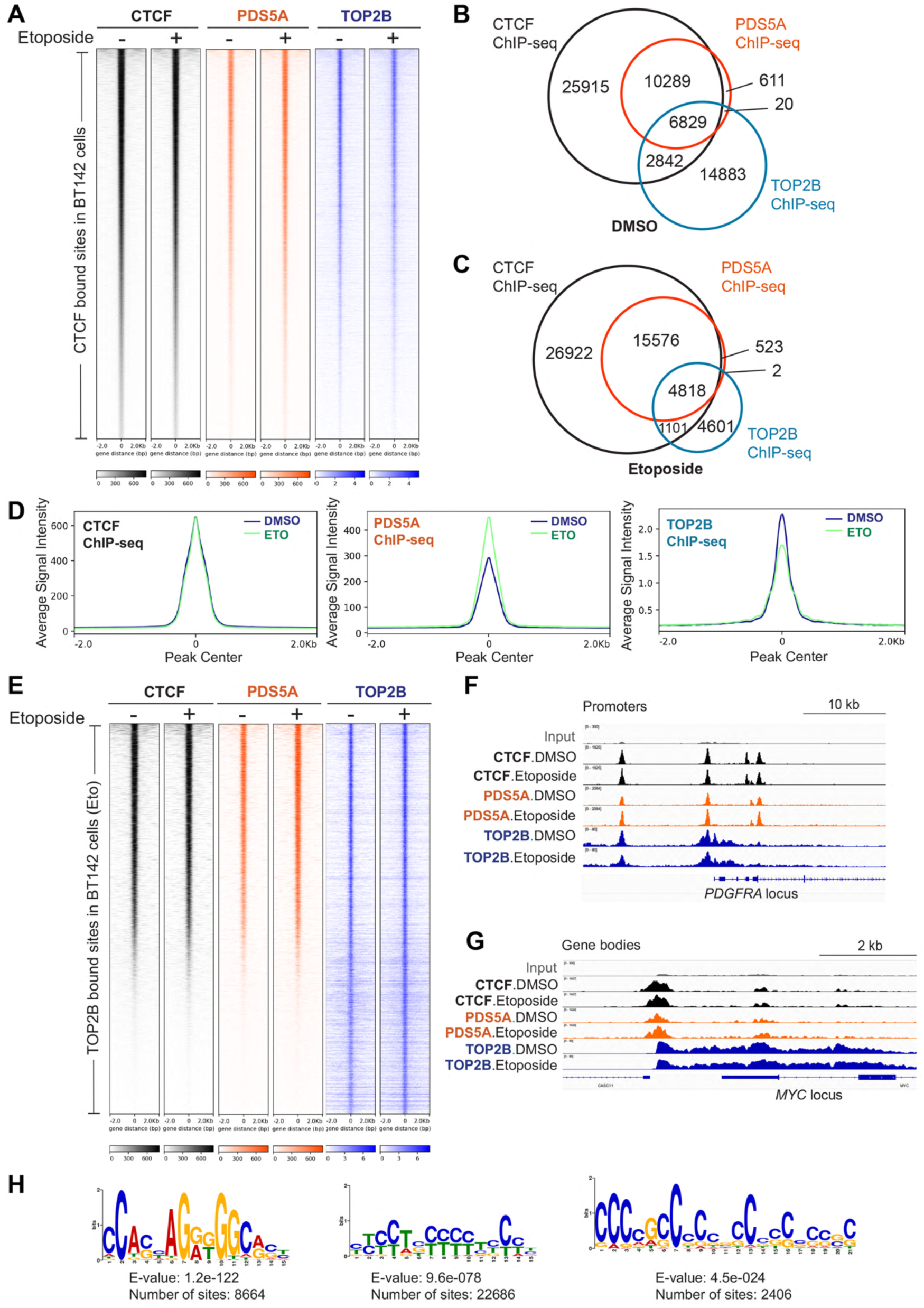
Gain of PDS5A is enhanced by TOP2B activity at CTCF binding sites in BT142 human glioma cells. **(A)** Heat maps showing CTCF, PDS5A, and TOP2B ChIP-seq read densities at CTCF-bound chromatin sites in BT142 cells under DMSO versus ETO treatment conditions. ChIP-seq data is from one representative of two biological replicates for CTCF and TOP2B, and one replicate for PDS5A. One replicate of TOP2B ChIP-seq has been re-analyzed^45^. **(B-C)** Venn diagrams showing CTCF, PDS5A, and TOP2B binding in BT142 under DMSO versus ETO treatment conditions. **(D)** Average density profile plots showing CTCF (left panel), PDS5A (middle panel), and TOP2B (right panel) ChIP-seq read densities under conditions of DMSO versus ETO treatment at CTCF-bound sites in BT142 cells. **(E)** Heat maps showing CTCF, PDS5A, and TOP2B ChIP-seq read densities at TOP2B-bound sites in BT142 cells under DMSO versus ETO treatment conditions. **(F-G)** Normalized ChIP-seq densities for CTCF, PDS5A, and TOP2B at *PDGFRA* (**F**) and *MYC* (**G**) loci under DMSO versus ETO treatment conditions in BT142 cells. These loci represent promoters (**F**) and gene bodies/intronic sites (**G**), respectively. **(H)** Motif analysis of TOP2B-binding regions in BT142 cells. Motifs associated with TOP2B peaks were identified by *de novo* MEME motif analysis, and the corresponding top matches (*i.e.* CTCF, MAZ, PATZ1, and VEZF1) are detailed in fig. S9 by Tomtom motif comparison tool. Motif search in MEME was performed *de novo* until 1000 sites were reached, and the corresponding e-values are depicted under the motifs.

In addition to their genome-wide co-occupancy, TOP2B and PDS5A specifically localized at highly expressed oncogenes, such as *PDGFRA* and *MYC* (Fig. 6F-G). To search for transcription factors that may be associated with the TOP2B binding sites in glioma cells, we used the TOP2B ChIP-seq data attained under DMSO conditions. We found that those TOP2B signals corresponded not only to CTCF binding motifs, but also to the binding motifs associated with other insulation factors including MAZ, PATZ1, VEZF1, and other ZNFs (Fig. 6H and fig. S9F), reflecting the results obtained above for PDS5A (fig. S3B). Overall, our results in gliomas highlight the existence of a previously unrecognized functional link between the active form of TOP2B and the enrichment of PDS5A on chromatin, mainly located at CTCF co-bound regions.

### PDS5A knockdown results in reduced occupancy of TOP2B on chromatin in BT142 glioma cells

Given the link between TOP2B catalytical activity and PDS5A enrichment on chromatin, we tested whether PDS5A functionally cooperates in localizing TOP2B on chromatin. To do so, we used three different shRNAs driven by an inducible TET-ON promoter in BT142 cells to knockdown (KD) PDS5A. TOP2B, PDS5A/B, and CTCF protein levels were not affected by induction of PDS5A KD with doxycycline (Dox), with histone H3 as loading control (Fig. 7A). Using this system, we assessed whether PDS5A KD affected TOP2B localization either at CTCF binding sites or at PDS5A binding sites genome-wide. We performed ChIP-seq in WT (untreated), and under KD (Dox) conditions, and observed that PDS5A KD resulted in the reduction of TOP2B signal at both CTCF-PDS5A-TOP2B overlapping sites (Fig. 7B, C, E, and see Fig. 7D, F as controls) and overall at PDS5A-bound sites genome-wide (fig. S10A, C, and see fig. S10B, D as controls). Global alterations in PDS5A, CTCF, and TOP2B binding were also noted upon PDS5A KD (see fig. S10E-K). As expected in the controls, PDS5A KD led to reduced PDS5A levels (Fig. 7C, D and fig. S10A-B and J-K), and did not affect CTCF localization (Fig. 7C, F and fig. S10A, D). Specifically, we observed that genes co-regulated by CTCF, PDS5A and TOP2B, such as *IDH2,* exhibited the loss of TOP2B upon PDS5A KD (Fig. 7G). Based on these results, we reasoned that either TOP2B was lost or re-localized on the genome upon PDS5A KD, compared to WT. Indeed, further analysis indicated that PDS5A KD appeared to lead to TOP2B re-localization in the genome (fig. S10G and fig. S11).

**Fig. 7.**
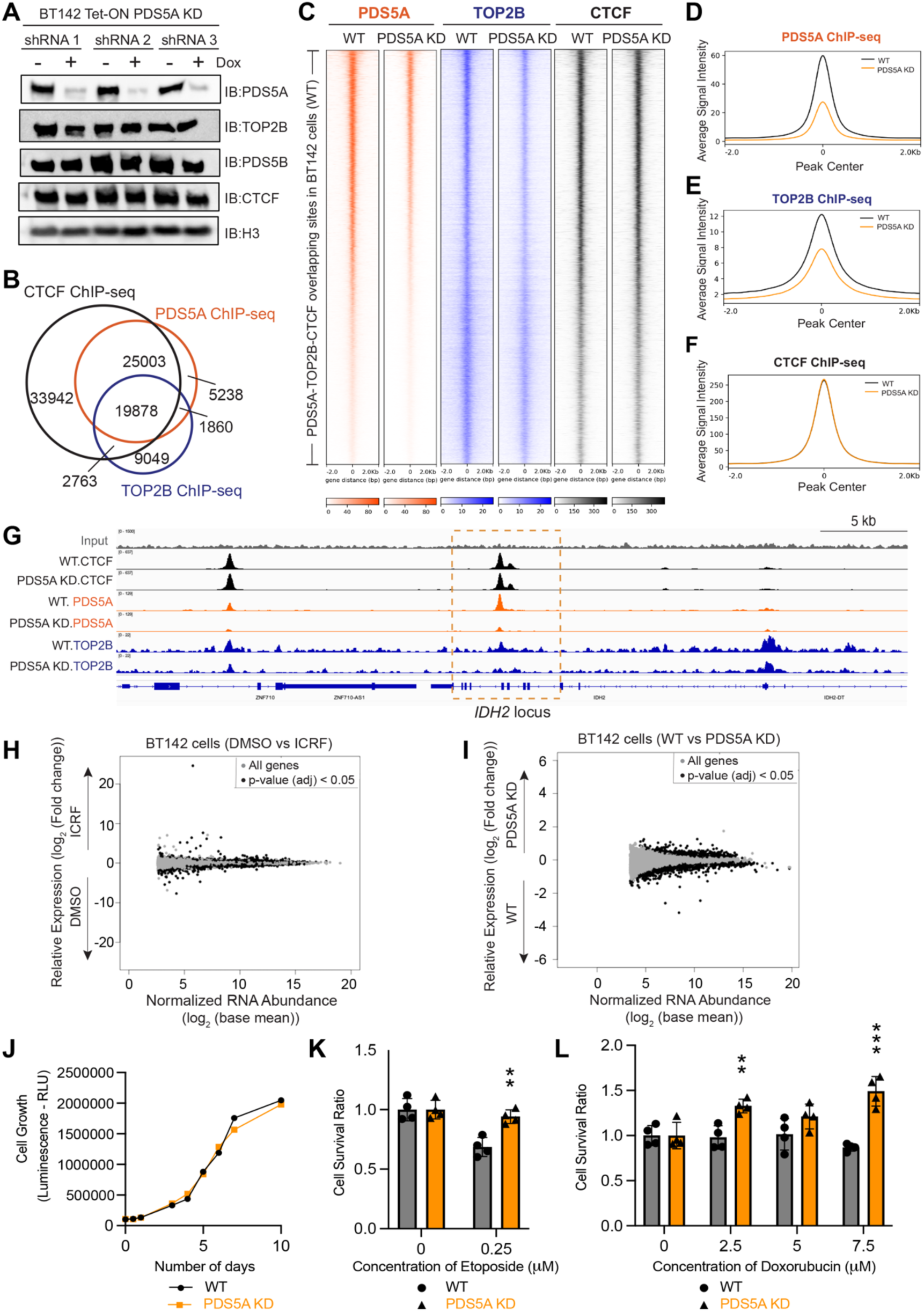
PDS5A knockdown (KD) results in reduced occupancy of TOP2B on chromatin, and PDS5A mediates sensitivity to TOP2-drugs in BT142 glioma cells. **(A)** BT142 cells were transfected with three different Tet-ON shPDS5A for inducible PDS5A KD as determined by WB of PDS5A, TOP2B, PDS5B, CTCF, and histone H3 (n=2). H3 serves as a loading control. IB: Immunoblot. **(B)** Venn diagram indicating the overlap of CTCF (black), PDS5A (orange), and TOP2B (blue) ChIP-seq binding sites in BT142 cells. **(C)** Heat maps showing PDS5A, TOP2B, and CTCF ChIP-seq read densities at PDS5A-TOP2B-CTCF common binding sites (n=19878) in BT142 cells in WT versus PDS5A KD conditions. **(D-F)** Average density profile plots showing PDS5A (**D**), TOP2B (**E**), and CTCF (**F**) ChIP-seq read densities at PDS5A-bound sites in BT142 cells in WT versus PDS5A KD conditions. **(G)** Normalized ChIP-seq read densities of CTCF, PDS5A, and TOP2B in WT versus PDS5A KD conditions at the *IDH2* locus, which is associated with tumor progression in BT142 cells. ChIP-seq data is from one representative biological replicate for each condition. **(H)** RNA-seq MA plot showing the impact of treatment with TOP2B inhibitor (ICRF-193, 15 μM, 6 hr) in BT142 glioma cells (TOP2B ICRF-193 inhibition re-analyzed data^45^). DMSO treatment serves as control. Gray dots indicate all genes, and black dots indicate DEGs across the conditions compared. RNA-seq data for control (DMSO) and ICRF-193 treated conditions are from five and seven biological replicates, respectively. **(I)** RNA-seq MA plot of BT142 cells carrying inducible Tet-ON shPDS5A under WT (-Dox) vs PDS5A KD (+Dox) conditions from four biological replicates. Gray dots indicate all genes and black dots indicate DEGs across the conditions compared. **(J)** The cell growth curves of WT vs PDS5A KD BT142 cells indicating that PDS5A KD did not affect cell growth. BT142 cells carrying Tet-ON shPDS5A, either unexpressed [WT (-Dox)] or expressed [PDS5A KD (+Dox)], were gauged for cell growth for 10 days in three biological replicates and showed no statistical differences. **(K-L)** PDS5A mediates sensitivity to TOP2-drugs. Shown are the surviving fraction of BT142 cells carrying Tet-ON shPDS5A, either unexpressed [WT (-Dox)] or expressed [PDS5A KD (+Dox)], and treated for 72 hr with TOP2-drugs in four biological replicates: (**K**) Etoposide (0 and 0.25 μM) or (**L**) Doxorubicin (0, 2.5, 5, and 7.5 μM). Two-sided Student’s t-test without multiple test correction was used (***p < 0.001, **p < 0.01).

To gain insight into those genes that may be co-regulated by PDS5A, TOP2B, and CTCF, we evaluated global gene expression using RNA-seq after the pharmacological inhibition of TOP2 for 6 hr with ICRF-193, an inhibitor of TOP2A/B enzymatic activity. The pharmacological inhibition of TOP2 resulted in ∼4000 genes (padj < 0.05) being differentially expressed as compared to the untreated case (Fig. 7H, see fig. S12A for gene ontology analysis). Given that TOP2B activity enriched PDS5A on chromatin in gliomas, we tested whether PDS5A KD affects gene expression similar to TOP2 enzymatic inhibition. Upon inducing PDS5A KD in BT142 glioma cells, ∼2600 genes (padj<0.05) were differentially expressed (Fig. 7I). Interestingly, we found that PDS5A KD affected biological pathways (fig. S12B), some of which resembled those affected by TOP2 enzymatic inhibition (fig. S12A). Taken together, these findings showed that catalytical inhibition of TOP2B and the reduction in PDS5A levels have an overlapping influence on gene expression and on the cellular processes impacted in glioma cells.

### PDS5A genome occupancy mediates sensitivity to TOP2 cancer drugs in gliomas

We next examined if the observed TOP2B re-localization on the genome upon PDS5A KD can mediate the sensitivity/resistance capacity of BT142 glioma cells to TOP2-drugs. First, we confirmed that the cell growth of BT142 cells was similar whether the Tet-ON shPDS5A was present under WT (-Dox) or PDS5A KD (+Dox) conditions (Fig. 7J). We then analyzed if PDS5A levels modulate resistance/sensitivity to TOP2-drugs by treating PDS5A-WT versus PDS5A KD BT142 cells with TOP2-targeting drugs (*i.e.* Etoposide or Doxorubicin). Upon treatment with Etoposide (0, 0.25 μM) (Fig. 7K) or Doxorubicin (0, 2.5, 5, 7.5 μM) (Fig. 7L), a fraction of PDS5A KD BT142 glioma cells survived as compared to the control, PDS5A-intact BT142 cells. Thus, modulation of PDS5A levels within glioma cells mediates their sensitivity to treatment with TOP2-drugs.

## Discussion

The three-dimensional chromatin architecture facilitates transcription, DNA replication, recombination, and chromatin remodeling. While these cellular processes alter the topological state of the genome, the topoisomerases actively keep the DNA in its proper state^62^. Specifically, TOP2B interacts with cohesin and CTCF on chromatin to release DNA tortional stress^39,40^. Although TOP2B-induced DSB resolves DNA topological problems during transcription^37,38^, DSB generate recurrent hotspots for chromosomal rearrangements that drive cancer^42,43^. Furthermore, TOP2B co-localization with CTCF and/or the cohesin complex (*i.e.* CTCF-RAD21-TOP2B and/or RAD21-TOP2B binding sites) promotes somatic mutations observed in many cancers^44^. Yet, the molecular mechanism of TOP2B recruitment has remained elusive. Here we show that PDS5A, a regulatory subunit of the cohesin complex, cooperates with TOP2B for their recruitment on chromatin in the context of CTCF and the cohesin complex in mouse cells, as well as in human glioma models.

We observed that PDS5A/B and TOP2B co-localized with CTCF, and that TOP2B activity or the product of its activity (*i.e.* DSB) enriched PDS5A, but not PDS5B, genome-wide. Furthermore, motif analyses of *de novo* PDS5A binding regions indicated motifs that match not only that of CTCF but also of other insulation factors (*i.e.* MAZ, PATZ1, and ZNFs), suggesting that the cooperative recruitment of PDS5A-TOP2B occurs at chromatin boundaries (Fig. 8A). Importantly, PDS5A remains bound on chromatin upon CTCF degradation. This outcome is consistent with a model in which other insulation factors identified in our motif analyses cooperate to recruit or maintain PDS5A at chromatin boundaries. Supporting this model, we previously observed that the loss of PDS5A leads to a further disruption of the *Hoxa5|6* boundary in cervical MNs that had been derived from mESCs bearing CTCF motif deletions at the *HoxA* cluster. In this differentiation system, the *Hoxa7* gene that had already been derepressed upon the loss of CTCF binding was now aberrantly upregulated upon the additional absence of PDS5A; accordingly, PDS5A was detected as a top-ranking candidate in CRISPR loss-of-function screens^14^. Overall, our results point to the unique enrichment of PDS5A genome-wide, including at chromatin boundaries as delimited by CTCF or other insulation factors, through TOP2B enzymatic activity that is critical for the resolution of topological stress in cells.

**Fig. 8.**
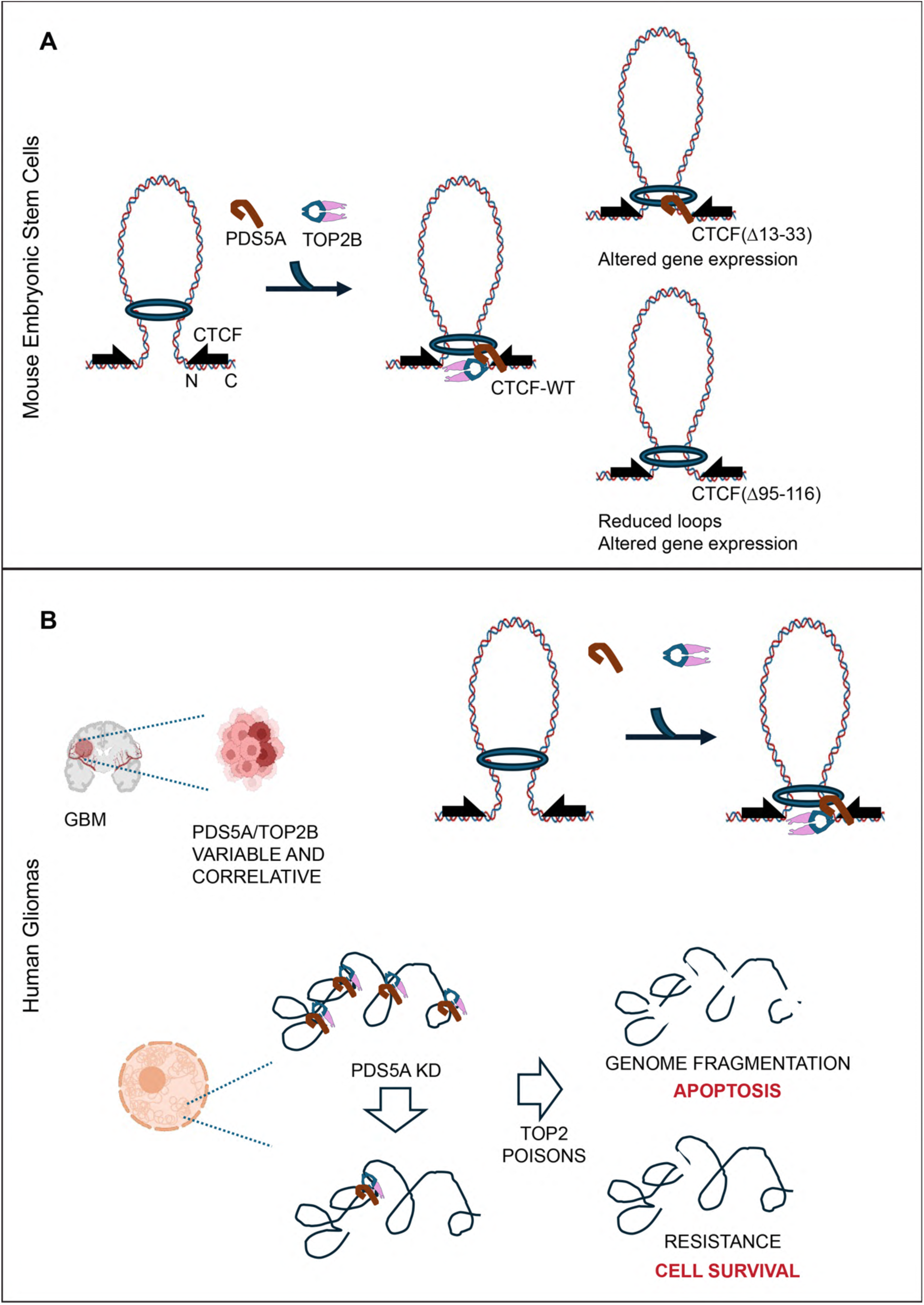
Model for PDS5A and TOP2B cooperation for chromatin recruitment. **(A)** In mESCs, PDS5A and TOP2B interact with CTCF, and TOP2B enzymatic activity or its product (*i.e.* DSB) enriches PDS5A on chromatin. An intact CTCF N-terminus is required for PDS5A-TOP2B interaction. While deletion of amino acids 13-33 at the N-terminus of CTCF, Δ13-33aa, affected TOP2B interaction, Δ95-116aa disturbed TOP2B-PDS5A interaction on chromatin, leading to altered gene expression and altered looping interactions in mESCs. **(B)** Consistent with this cooperation between PDS5A and TOP2B, their individual expression levels in gliomas are variable and correlative. Moreover, PDS5A-TOP2B are functionally linked in gliomas. An inducible PDS5A KD directly impacts TOP2B localization leading to its reduced chromatin occupancy. Partial arrows (black) on the DNA indicate convergent CTCF DNA-binding sites required for establishing CTCF-mediated chromatin anchors. The circle (blue) represents the cohesin complex as it extrudes a chromatin loop. The levels of PDS5A promote glioma sensitivity or resistance to treatment with TOP2-drugs. Created with BioRender.com.

In addition to a previously reported CTCF-PDS5A interacting region^17^, we identified a novel CTCF-PDS5A interacting region that facilitates TOP2B localization at chromatin boundaries. Notably, deletions in these CTCF-PDS5A interacting regions affected different gene sets, validating that these regions are not redundant. In particular, the deletion of CTCF-PDS5A interaction region, CTCF-MUT2(Δ95-116), led to differentially expressed genes that overlap with PDS5A binding sites, and reduced looping interactions in mESCs. Underscoring the cooperation between PDS5A-TOP2B, the loss of either of the CTCF-PDS5A interacting regions reduced TOP2B interaction upon Etoposide-treatment, known to trap active TOP2B. In accordance, PDS5A knockdown resulted in decreased TOP2B occupancy at CTCF-PDS5A-TOP2B occupied regions in glioma cells. Thus, we propose that proper gene expression and genome organization require an intact N-terminal domain of CTCF that is critical for its interaction with PDS5A-TOP2B, and that this interaction is modulated upon activating TOP2B.

Cell-type-specific distal gene regulation is dependent on the cohesin loading cycle, including its accessory loading and unloading activities^20,63,64^. Previous work showed that PDS5A/B^16,18^, WAPL^16^, and cohesin acetylation control chromatin loop length at CTCF binding sites^63,65–68^, consistent with the reduction of loops in CTCF-MUT2(Δ95-116) cells. The unloading of the cohesin complex from chromatin is facilitated by the PDS5-WAPL interaction^52,69,70^, yet our study did not detect WAPL interaction with PDS5A-TOP2B, nor WAPL enrichment on chromatin by TOP2B activity. In relation to cohesin loading, whereas the interaction of NIPBL-cohesin favors cohesin ATP-ase activity^71^, the interaction of PDS5A-cohesin results in a reduction of this cohesin ATP-ase activity^20,65,71^. However, the molecular mechanism of the exchange between NIPBL and PDS5A on cohesin at chromatin boundaries is not understood. Further biochemical experiments could address whether TOP2B activity favors PDS5A exchange for NIPBL on cohesin at CTCF binding sites or other insulation factor binding sites. Indeed, Pds5 is involved in the inhibition of loop enlargement despite the lack of CTCF in yeast^67,72^, and overexpression of Top2 specifically suppressed temperature sensitivity of Pds5 mutants in yeast^73^. Given that the activity of TOP2B regulates PDS5A chromatin occupancy and that knockdown of PDS5A alters TOP2B occupancy reciprocally, our study points to an interplay between PDS5A-TOP2B and CTCF in cohesin dynamics and the regulation of loops.

In GBM, some genetic alterations have been associated with epigenetic mechanisms such as DNA methylator phenotype (CIMP) in *IDH1/2* mutant cells^74,75^, insulator dysfunction^29^, histone mutations of H3K27^76–78^, and three-dimensional genome organization^79–82^. Based on our results indicating the interaction of PDS5A-TOP2B-CTCF, the dependence of PDS5A occupancy on TOP2B enzymatic activity in mESCs and gliomas, and the previously shown link between TOP2B enzymatic activity and oncogene expression in gliomas^45^, we further investigated the correlation of TOP2B and PDS5A expression in gliomas. Given that PDS5A and TOP2B cooperate for their recruitment on chromatin via CTCF, perturbing the interaction between CTCF-PDS5A impacted the number of loops in mESCs and the localization of TOP2B in both mESCs and glioma cells. Interestingly, reduced TOP2B occupancy is observed upon PDS5A knockdown while the enrichment of PDS5A on chromatin required TOP2B enzymatic activity. Consistent with the model proposed (Fig. 8B), PDS5A knockdown in glioma cells leads to a relocation of TOP2B on chromatin that further contributes to the heterogeneity in gene expression reported and thus, might mediate variable responses to TOP2 drugs.

Many cancer drugs targeting TOP2A/B, such as Etoposide and Doxorubicin, are frequently used in cancer therapies. However, secondary malignancies have been associated with TOP2A/B at CTCF binding sites^42^. In addition, CTCF became a hotspot frequently associated with some translocations in therapy-related myeloid leukemias (t-AML) and cancer drug resistance^42,50^. As TOP2B contributes to the disentanglement of CTCF/other insulator-mediated chromatin loops to release torsional stress affecting vital cellular processes, TOP2B enzymatic function is directly implicated in TOP2-drug resistance in cancer^83,84^. Strikingly, TOP2B relocation upon PDS5A knockdown facilitates resistance to Etoposide and Doxorubicin in glioma cells. Thus, we envision that the dynamics of PDS5A-TOP2B occupancy in the genome might clarify GBM-associated heterogeneity in expression and importantly, inform about sensitivity to TOP2-drug response. Altogether, our study highlights the cooperativity of PDS5A-TOP2B for their genomic localization and demonstrates that PDS5A mediates sensitivity to TOP2-drugs in glioma cells.

### Limitations of the study

The TOP2 inhibitors used in this study induce both TOP2A and TOP2B cleavage complexes. Yet, we present complementary gene silencing experiments supporting that PDS5A interacts specifically with TOP2B in glioma cells. We also studied PDS5A and TOP2B interaction in mESCs as a representative normal phenotype to support our conclusions. We further addressed this limitation by presenting experiments in MNs differentiated from mESCs. The TOP2B protein levels are higher in MNs relative to the undifferentiated cells, and TOP2A protein levels decreased upon differentiation. It is worth noting that we relied on glioma cell lines in culture, in which characterization of gene expression and ChIP-seq were performed *in vitro*. Such *in vitro* experiments could be influenced by artefacts and changes in tumor cell phenotype that relate to culture conditions. These latter limitations were addressed by our complementary analysis of human glioma and brain tissues.

### Materials and Methods Cell Culture

E14 mESC (ES-E14TG2a) cell line from ATCC (CRL-1821), was cultured in standard medium supplemented with LIF, and 2i conditions (1 mM MEK1/2 inhibitor (PD0325901, Stemgent) and 3 mM GSK3 inhibitor (CHIR99021, Stemgent)). Differentiation into cervical motor neurons (MNs) was performed using a protocol described previously^14,15^. Briefly, mESCs were differentiated into embryoid bodies (EBs) in 2 days, and further patterning was induced with the addition of 1 μM all-trans-retinoic acid (RA, MilliporeSigma) and 0.5 μM smoothened agonist (SAG, Calbiochem). 293FT cells were obtained from Thermo Fisher Scientific (R70007) and cultured in standard medium as described in the manufacturer’s protocol. The identity of the BT142 IDH1 R132H cell line from ATCC (ACS-1018) was confirmed by short tandem repeat (STR) and the cells were cultured with NeuroCult NS-A Proliferation kit (Catalog No. 5751, Stem Cell Technologies), 20 ng/mL recombinant human Epidermal Growth Factor (EGF, Catalog No. 100-15, PeproTech), 100 ng/mL recombinant human Platelet-Derived Growth Factor-AA (PDGF-AA, Catalog No. 100-13A, PeproTech), 20 ng/mL recombinant human Fibroblast Growth Factor (R&D Systems, Catalog No. 233-FB), 2 µg/mL heparan sulfate (Catalog No. H3149, Sigma), and antibiotics (1X)^45^.

### Chromatin Immunoprecipitation and Sequencing (ChIP-seq) and Native ChIP-seq under ETO/DMSO Conditions

ChIP-seq-ETO and native ChIP-seq were performed with 5×10^7^ cells that were harvested after growth in cell suspension. For ChIP-seq, the cells were treated with 500 μM Etoposide or DMSO for 10 min in culture media, washed 1X with PBS, and then resuspended in crosslinking media using 1% formaldehyde (1% FA) or non-FA, as indicated. Nuclear extracts were obtained, and chromatin was fragmented to ∼500 bp using a Covaris sonicator. Ten micrograms of the antibodies listed in Table S1 were incubated overnight. In addition, chromatin from *Drosophila* (1:100 ratio to mESC-derived or BT142-derived chromatin), and *Drosophila*-specific H2Av antibody were used as a spike-in control in each sample. Then, washes were performed with RIPA buffer four times followed by a final wash of TE containing 50 mmol/L NaCl. The crosslink was reversed, and DNA was purified using phenol:chloroform:isoamyl alcohol for library preparation and sequencing.

### Immunoprecipitation (IP)

Nuclear extracts were prepared as described previously^85,86^. Briefly, buffer A (10 mM HEPES, pH 7.9, 1.5 mM MgCl_2_, 10 mM KC1 and 0.5 mM DTT) was used to remove the cytoplasmic fraction, and buffer C (20 mM HEPES, pH 7.9), 25% (v/v) glycerol, 420 mM NaCl, 1.5 mM MgCl_2_, and 0.2 mM EDTA) was used to extract nuclear proteins. Anti-FLAG M2 Magnetic Beads (Millipore Sigma, #M8823) were used for IP, and proteins bound to the beads were released by competitive elution using 3X FLAG peptide. For other IP experiments, the specific antibodies used are listed in Table S1.

### Expression of Recombinant Proteins

cDNA of the protein of interest was cloned into CβF plasmid and transfected into 5×10^8^ 293FT cells using Polyethylenimine (PEI). After 60 hr, cells were harvested by washing with cold 1X PBS, scraping and incubating with TMSD buffer (20 mM HEPES, 5 mM MgCl_2_, 85.5 g/L sucrose, 25 mM NaCl, 1 mM DTT, and protease inhibitors) for 10 min. After centrifugation to pellet the cells, the supernatant containing the cytoplasmic fraction was removed. The nuclei fraction was further incubated in BA450 buffer (20 mM HEPES, 450 mM NaCl, 5% glycerol and protease inhibitors) for 30 min, the insoluble material was pelleted, and the supernatant containing the nuclear extract was used for FLAG affinity purification for 12 hr. FLAG beads were washed 3X with BA450, and then 3X with BA50 (20 mM HEPES, 50 mM NaCl, 5% glycerol). Ultimately, the proteins were eluted using 3X FLAG peptide with rotation at 4°C overnight.

### IP of FLAG-mPDS5A1-750aa

CβF-mPDS5A1-750aa construct was transfected into 293FT cells using PEI, and nuclei were fractionated using TMSD buffer (20 mM HEPES, 5 mM MgCl_2_, 85.5 g/L sucrose, 25 mM NaCl, 1 mM DTT, and protease inhibitors). After cytosol removal, nuclear extraction was processed as described above for recombinant proteins using BA450 buffer (20 mM HEPES, 450 mM NaCl, 5% glycerol and protease inhibitors). FLAG affinity purification and 3X FLAG peptide elution was performed in BA50 buffer with rotation at 4°C overnight.

### RNA-sequencing

Total RNA from mESCs or BT142 cells was purified with RNAeasy Plus Mini kit (Qiagen) using the manufacturer’s instructions. 1 μg RNA was used to prepare standard RNA-seq libraries according to the manufacturer’s instructions by Onco-Genomics Shared Facilities at the University of Miami Miller School of Medicine.

### Western Blot (WB) Analysis

Cells were washed in 1X PBS and resuspended in RIPA buffer (150mM NaCl, 10mM HEPES, pH 7.9, 1mM EDTA, 1% Triton X-100, 0.1% SDS, 0.1% Sodium deoxycholate), incubated for 20 min, and proteins were spun-down and quantified. WB analysis was used to evaluate protein abundance using the antibodies. Antibodies used are listed in Table S1.

### Electrophoretic Mobility Shift Assays

Single-stranded 5’Cy3-oligonucleotides containing the cHS4 FII CTCF binding site were resuspended in a 100 mM stock solution using 10 mM HEPES (pH 7.5) and annealed. 1mM probe was resuspended in binding buffer (10 mM HEPES, pH 7.5, 50 mM NaCl, 5% glycerol, 5 mM MgCl_2_, 0.1 mM ZnSO4, and 0.1 μg salmon sperm DNA). The reactions were then incubated with increasing amounts (0.05, 0.1, 0.2 μg) of mouse recombinant CTCF, either WT or CTCFMut2(Δ95-116) for 1 hr, 25°C and/or PDS5A (0.4 μg) for 1 hr at 25°C. The reactions were then run on 5.2% acrylamide gels for 30 min at rt with 150 V and 0.25X TBE buffer. Finally, the acrylamide gel was exposed using the imager settings for UV-Cy3.

### Immunohistochemistry (IHC)

Formalin-fixed, paraffin-embedded human glioma tissue microarray (TissueArray.com GL2082a) was deparaffinized with xylene and antigen retrieval was performed with 10 mmol/L sodium citrate buffer (pH = 6). IHC staining was performed on a Leica Bond-Max automatic immunostainer. IHC was performed for PDS5A (Novogen), PDGFRA (Cell Signaling Technology, 5241) and TOP2B (Abcam, ab72334) antibodies. Counterstaining was done with hematoxylin and eosin. Visiopharm software was used to quantify the PDS5A-, PDGFRA-, or TOP2B-positive cells per total cell number as indicated in the figure legend. Parameters were adjusted to detect nuclei first by hematoxylin staining. We applied a threshold to the mean intensity of DAB chromogen for all the cells to define positive staining nuclei for PDS5A and TOP2B. Cytoplasmic PDGFRA staining was quantified after applying a threshold to the mean intensity of the DAB chromogen for all cells. The same thresholds and parameters were used for all tumor samples.

### CTCF WT and Mutant Rescues

1×10^5^ CTCF-degron (CTCF-GFP-AID-Tir1) cells were transfected using Lipofectamine reagent (Invitrogen, 11668019) with pPB-CAG-FLAG-HA-CTCF WT, pPB-CAG-FLAG-HA-CTCFΔ13-33aa, or PB-CAG-FLAG-HA-CTCFΔ95-116aa. After 72hr, cells were selected with Hygromycin 0.2mg/mL and protein expression levels were assessed by WB.

### shRNA-PDS5A TET-ON system in BT142 cells

BT142 cells were infected with shRNA-PDS5A-TET-ON system viral particles for 12 hr. Cells were selected with 1μg/mL Puromycin after 72 hr post infection and protein expression levels were tested. PDS5A knockdown following 1μg/mL Doxycycline for 72 hr and the shRNA-based PDS5A silencing in a TET-ON system was determined by WB.

### Micro-C Preparation

Dovetail® Micro-C Kit was utilized for Micro-C library preparation according to the manufacturer’s protocol. 1 million cells were used for Micro-C experiments in each replicate in WT versus CTCF-MUT2(Δ95-116) conditions across duplicates. Briefly, disuccinimidyl glutarate (DSG) and formaldehyde was used for crosslinking chromatin in the nucleus. Then, the crosslinked chromatin was digested with micrococcal nuclease (MNase) *in situ*. After digestion, chromatin fragments were extracted by lysing with SDS, and binding to Chromatin Capture Beads. Next, chromatin ends were repaired and ligated to a biotinylated bridge adapter through proximity ligation. Finally, the crosslinks were reversed, the associated proteins were degraded, and the DNA was purified and used for library preparation using Illumina-compatible adaptors. After library preparation, biotin-containing fragments were isolated using streptavidin beads before PCR amplification. Micro-C libraries were sequenced on an Illumina Novaseq X platform to generate > 800 million 2 x 150 bp read pairs that ensures the required data depth for loop analysis (see Table S2 for the number of reads across Micro-C samples).

### Cell Growth Curve

BT142 cells expressing Tet-ON shPDS5A under WT (-Dox) vs PDS5A KD (+Dox) conditions were seeded onto 96-well plates at a concentration of 2000 cells/well. Doxycycline was added at day 0.5, and the cells were followed for 10 days. CellTiter-Glo 2.0 (G9242) luminescent reagent was used for lysis and detection of viable cells. Three technical replicates per condition in two independent experiments were performed. A t-test showing P<0.05 was considered significant.

### Cell Survival Assay

BT142 cells expressing Tet-ON shPDS5A under WT (-Dox) vs PDS5A KD (+Dox) conditions were seeded onto 96-well plates at a concentration of 5000 cells/well. TOP2 drugs, Etoposide (Tocris, 1226) (0 and 0.25 μM), and Doxorubicin (Sigma-Aldrich, D1515) (0, 2.5, 5, and 7.5 μM) were added to the media. After 0, 24, 48, and 72 hr of treatment, CellTiter-Glo 2.0 (Promega, G9242) luminescent reagent was used for lysis and detection of viable cells. Four technical replicates per condition in two independent experiments were performed. A t-test showing P<0.05 was considered significant.

### Data Analysis ChIP-seq analysis

ChIP-seq data was analyzed as described previously^14,15^. Briefly, Bowtie 2 (version 2.3.4.1) was used for mapping the sequence reads to mm10, hg38, or dm6 reference genomes using default parameters^87^. Quality filtering and duplicate removal were performed by using SAMtools (version 1.9)^88^. MACS (version 1.4.2) was used for narrow peak calling using default parameters of ‘macs2’^89^. After normalization with spike-in read counts or total read counts as indicated, ChIP-seq heat maps, density plots, venn diagrams and tracks were generated. DeepTools in R was used for generating heat maps^90^. Venn diagrams were generated by using ‘ChIPpeakAnno’ package (version 3.36.1) from Bioconductor^91^. The Venn diagram sizes were drawn using the online tools, “http://barc.wi.mit.edu/tools/venn/” and “https://www.meta-chart.com/venn#/display”. Normalized ChIP-seq tracks were visualized in Integrative Genomics Viewer (IGV, version 2.16.2)^92^. ChIP-seq replicates were evaluated by visualizing at Integrative Genomics Viewer and generating heat maps. PAVIS tools were used for ChIP-seq data annotation to the genes^93^. For motif analysis, ChIP-seq “bed” file coordinates were converted into “fasta” by using fetch sequences tool within Regulatory Sequence Analysis Tools (RSAT)^94^ and MEME (version 5.5.8) was used for motif analysis of PDS5A and TOP2B in mESCs and/or BT142 cells^95^, and Tomtom (version 5.5.8) was used as a motif comparison tool^96^. Motif search in MEME was performed *de novo* until 1000 sites were reached and the e-values corresponding to the motifs were depicted.

### Micro-C analysis

The Micro-C analysis was performed by Dovetail® using the tools shown in https://micro-c.readthedocs.io/en/latest/index.html. In brief, HiC contact matrices were generated in each sample by aligning fastq files and generating valid pairs bam files. After assessment of each replicate and merging the replicates, the loops were identified using HiCCUPs from the Juicer package^97^, and Mustache^98^, with default parameters. Micro-C comparative analysis between CTCF-WT versus CTCF-MUT2(Δ95-116) samples were performed using HiCcompare algorithm by Dovetail®^99^. As HiCcompare method applies a joint normalization for CTCF-WT versus CTCF-MUT2(Δ95-116) conditions before the differential analysis, this process ensures the statistical robustness of the loops called. In this method, an MD normalization plot was generated to visualize the effect of normalization between the conditions to ensure that the data was appropriate for the downstream difference detection. Moreover, the standard outputs generated here include a table of differentially interacting genomic coordinates, the frequency of interactions, the difference and the permutation p-values (for pipeline, see https://micro-c.readthedocs.io/en/latest/microc_compare.html). FAN-C toolkit^100^ was further used for generating AB compartments by Dovetail®, and the resulting tables were presented as bar plots in GraphPad Prism. APA analysis and visualization of loops: normalized APA analysis was performed for all the loops identified by Mustache (16 kb loops) in WT versus CTCF-MUT2(Δ95-116) ESCs using 10 kb resolution by using Juicer APA tools^101^. Visualization of the zoomed-in coordinates/loci in Micro-C was performed using Juicebox^97^. HiCcompare differential loop outcomes were plotted using Rstudio. Analysis of PDS5A, CTCF, RAD21 ChIP-seq occupancy at loop anchors: chromatin loops called by Mustache were used for overlapping with PDS5A, CTCF, and RAD21 ChIP-seq peaks by using pairToBed from the BEDTools suite (version 2.27.1)^102^ and pgltools (version 1.2.1)^103^. Afterwards, APA plots were generated at the overlapping regions using Juicer APA tools.

### RNA-seq analysis

RNA-seq data was analyzed as described for BT142 cells^45^ and for mESCs^14^. Briefly, the sequence reads were mapped to mm10 and hg38 reference genome using Bowtie 2 (version 2.3.4.1)^87^, and normalized differential gene expression was obtained with DEseq2 (version 1.26.0)^104,105^ by using the Wald test built into DEseq2 with an FDR cutoff of 0.05. Differential gene expression values and corresponding p-values are listed in Table S3, S5, S7, and S9. For Gene Ontology (GO) analysis, the PANTHER database was used^106^ and the relevant GO terms are listed in Table S4, S6, S8, and S10. Venn diagrams were drawn using online tools: GeneVenn (https://www.bioinformatics.org/gvenn/), Venn Diagram Generator (http://barc.wi.mit.edu/tools/venn/), and Venn Diagram Maker (https://www.meta-chart.com/venn#/display). In the case of re-analysis of RNA-seq upon TOP2 inhibition in BT142 cells^45^, the deposited gene counts in GEO (GSE133561) dataset were used for further analysis, as described above.

## Quantification and Statistical Analysis

Statistical analysis related to experiments have been described above in each section. In addition, the two-sided t-test was generally used in the cases where two populations were compared.

## Acknowledgments

We are grateful to Lynne Vales for the invaluable scientific discussions and guidance on the manuscript. We thank the past and present members of the Reinberg Laboratory for the scientific discussions. We thank the NYU Grossman School of Medicine’s Genome Technology Center for help with sequencing; the Applied Bioinformatics Laboratories for providing bioinformatics support. This study used the computing resources at the High-Performance Computing Facility of the Center for Health Informatics and Bioinformatics at the NYU Grossman School of Medicine. We thank the Miller School of Medicine’s Pathology Research Resources at the University of Miami. We also thank the Onco-Genomics Shared Resource (OGSR) of the Sylvester Comprehensive Cancer Center at the University of Miami, RRID: SCR_022502, for help with sequencing at the later stages of this work. We are deeply grateful to Francisco Javier Gonzalez Muñoz for the invaluable scientific discussions and advice.

## Author contributions

Conceptualization: EG-B, DR

Methodology: EG-B

Investigation: EG-B, HO

Data Analysis: HO, EG-B

Supervision: DR, EG-B

Writing—original draft: EG-B

Writing—review & editing: EG-B, DR, HO

## Data and materials availability

Further information and requests for resources and reagents should be directed to and will be fulfilled by the lead contact, Danny Reinberg. All unique/stable materials generated in this study are available from the corresponding author upon reasonable request with a completed Materials Transfer Agreement. Sequencing data has been deposited at Gene Expression Omnibus (GEO) and accession (GSE311476) will be available immediately upon publication. Publicly available dataset (GSE133561) was used for TOP2B ChIP-seq data in Fig. 6, and RNA-seq data in Fig. 7H and fig. S12A. Publicly available datasets (GSE157139 and GSE230482) were used for MAZ ChIP-seq and PATZ1 ChIP-seq in fig S2, respectively. WB raw data shown in the figures will be available in Mendeley: doi: 10.17632/kkvxyv5nrx.1. Code availability: Analysis tools used in this study have been published previously, as described in the Methods. The codes will be available upon reasonable request.

## Supplementary Materials

**Fig. S1.**
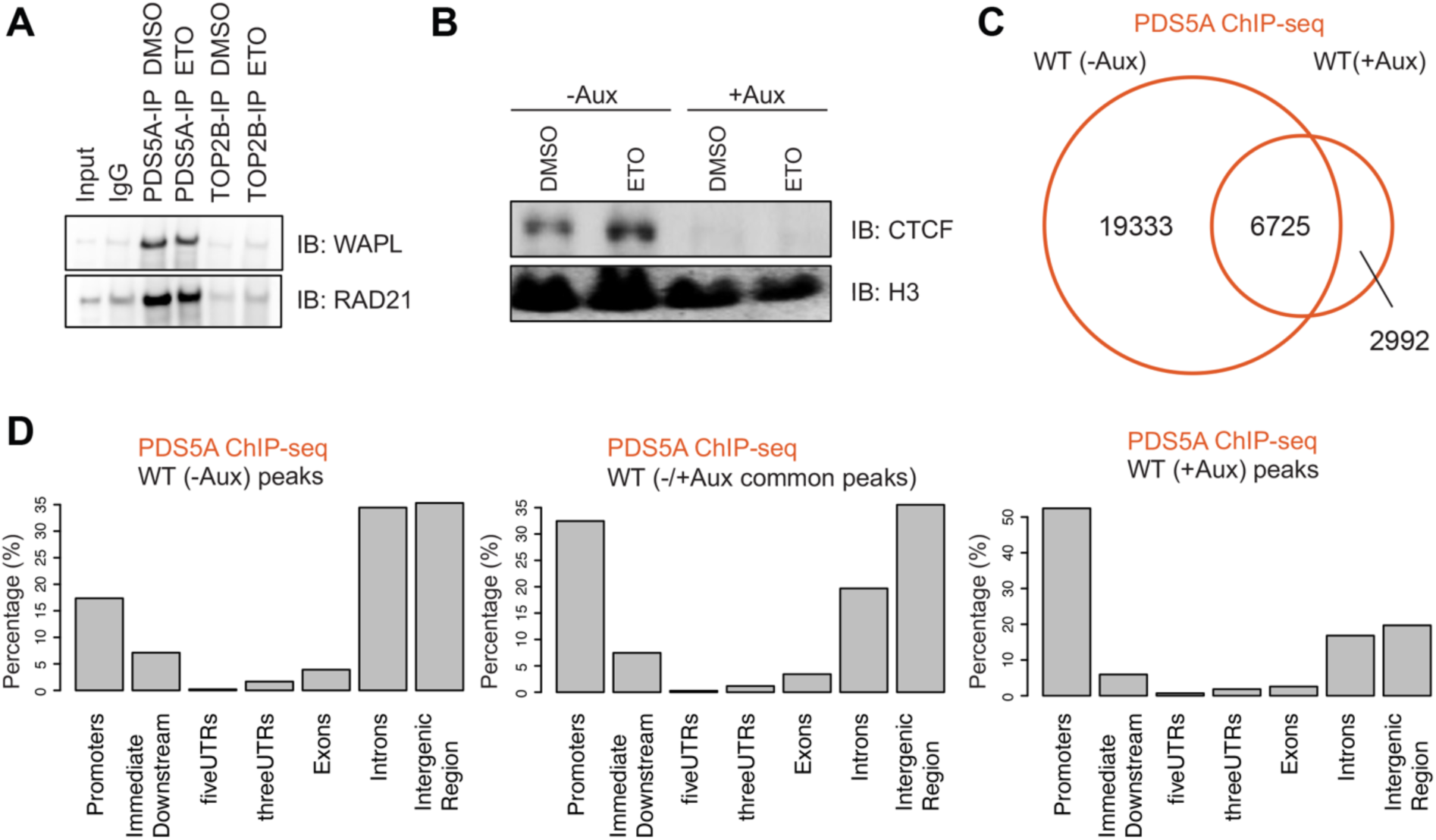
The activity of TOP2B enriches PDS5A on chromatin, related to Fig. 1-2. **(A)** PDS5A and TOP2B Immunoprecipitation (IP), in DMSO versus Etoposide (ETO) treatment conditions, and Western blot of WAPL and RAD21 in MNs (MN chromatin pellet treated with Benzonase nuclease). **(B)** CTCF Chromatin Immunoprecipitation (ChIP) and Western blot analysis in CTCF degron mESCs with CTCF intact (-Aux) or CTCF degraded (+Aux) in DMSO versus Etoposide (ETO) treatment conditions (two independent immunoprecipitations of 1% formaldehyde crosslinked chromatin experiments for CTCF ChIP). H3 serves as a loading control. **(C)** Venn diagram showing PDS5A binding (orange) in CTCF degron mESCs with CTCF intact (WT, -Aux) versus CTCF degraded (ΔCTCF, +Aux) conditions. **(D)** Bar plots indicating the distributions of genomic features for PDS5A binding regions at each part of the Venn diagram in (**C**). ChIP-seq data is from one representative of three biological replicates.

**Fig. S2.**
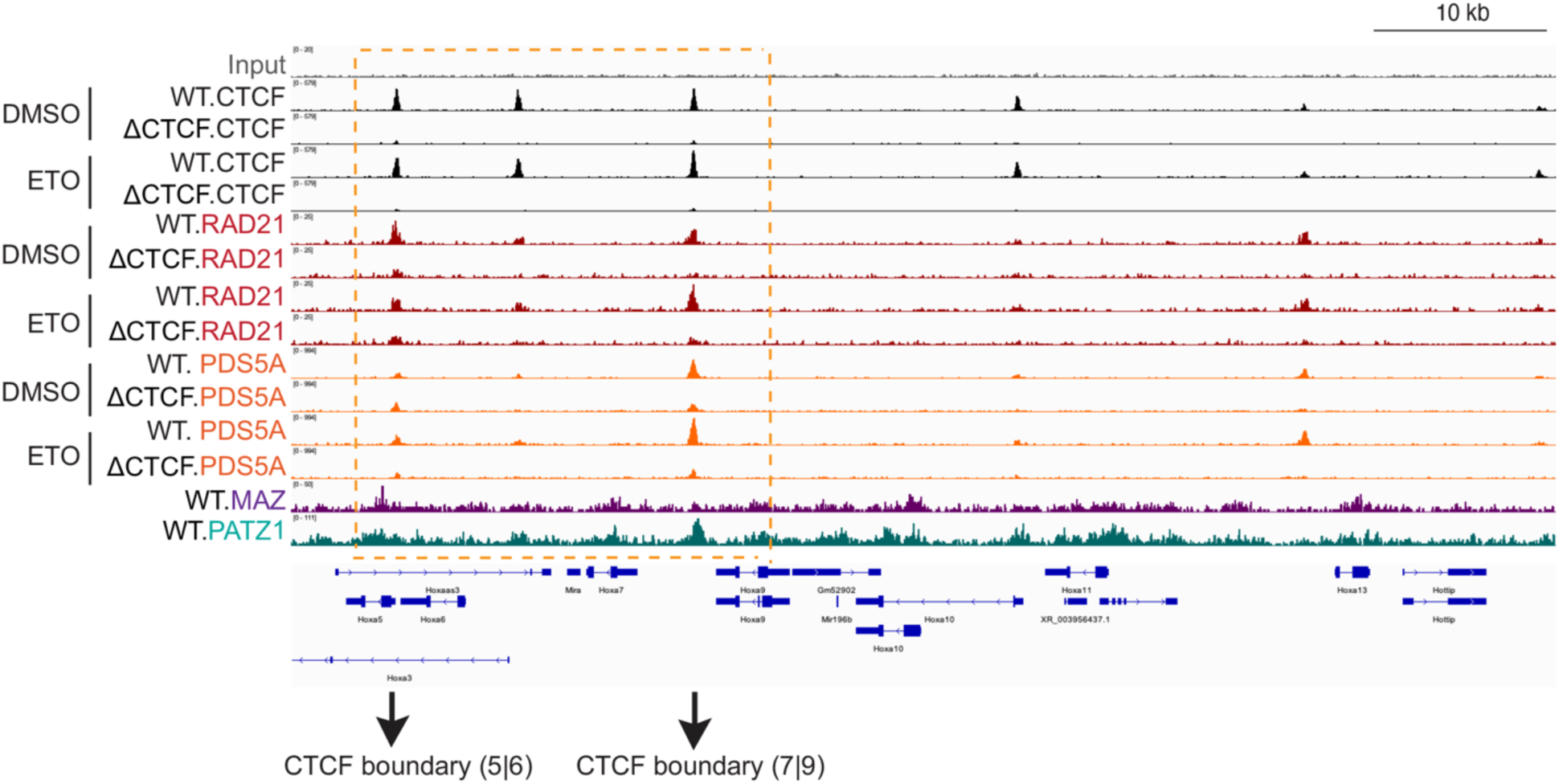
PDS5A remains bound on chromatin in the absence of CTCF in mESCs, related to Fig. 2. Normalized ChIP-seq densities for CTCF, RAD21, PDS5A, MAZ and PATZ1 at indicated regions (*marked in orange*) at *HoxA* cluster in CTCF intact (WT, -Aux) versus CTCF degraded (ΔCTCF, +Aux) mESCs in DMSO versus Etoposide (ETO) conditions. ChIP-seq data is from one representative of three biological replicates for PDS5A, two biological replicates for RAD21, and one replicate for CTCF across the conditions. ChIP-seq data for MAZ and PATZ1 are from the publicly available datasets (GSE157139 and GSE230482, respectively).

**Fig. S3.**
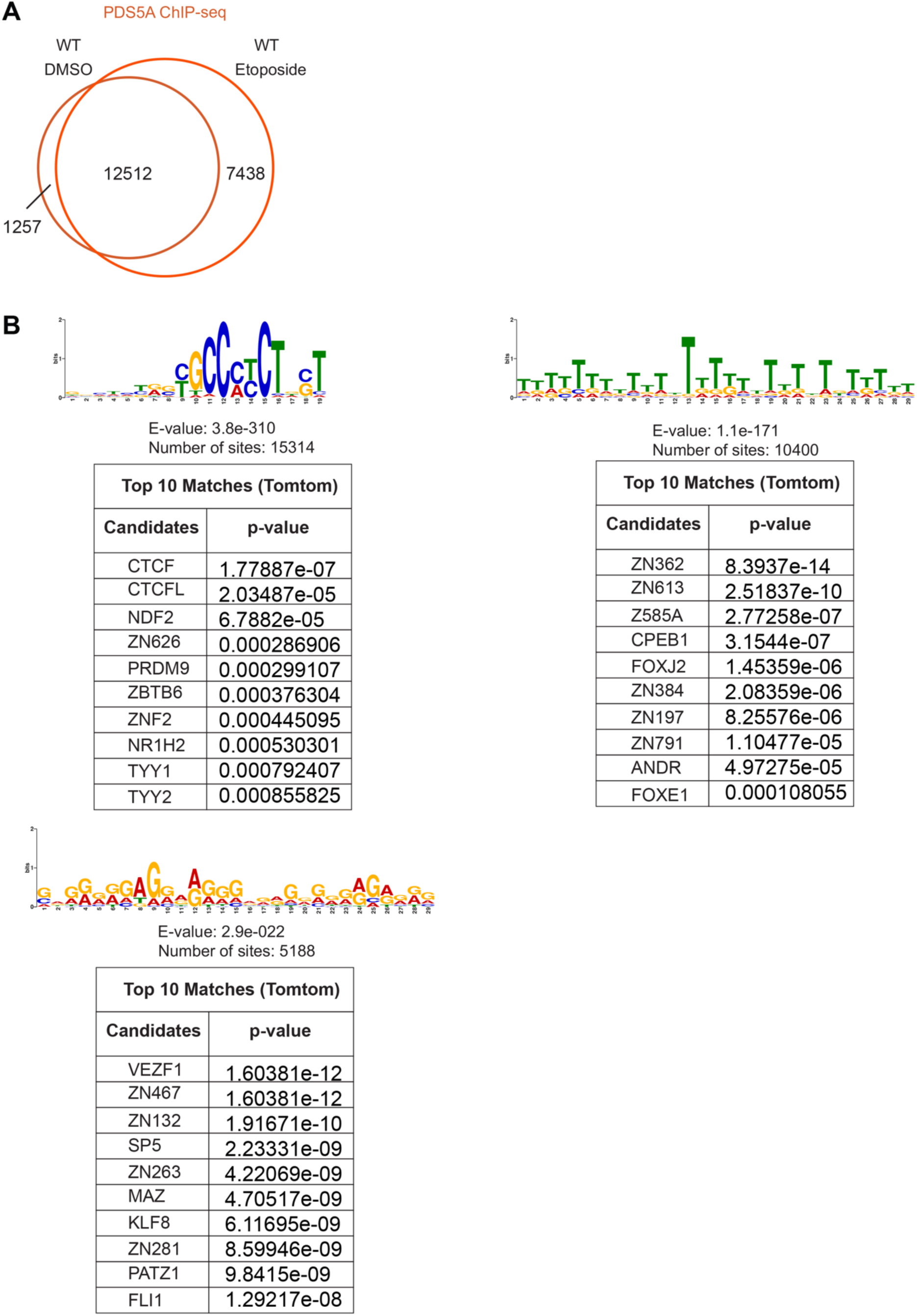
Gain of PDS5A is enhanced by TOP2 activity specifically at CTCF binding sites and other insulator factor motifs, such as MAZ, PATZ1, VEZF1, and ZN263, related to. Fig. 2**. (A)** Venn diagram showing PDS5A binding under conditions of DMSO versus Etoposide treatment in CTCF intact (WT, untreated) mESCs. **(B)** Motif analysis of PDS5A binding regions in ETO condition in mESCs (n=19,950 peaks). Motifs associated with PDS5A peaks were identified by *de novo* MEME motif analysis, and the corresponding top matches to the motifs were shown in the tables by using Tomtom motif comparison tool. Motif search in MEME was performed *de novo* until 1000 sites were reached, and the corresponding e-values to the motifs were depicted under the motifs and the corresponding p-values to the motif matches were depicted in the tables. ChIP-seq data is from one representative of three biological replicates.

**Fig. S4.**
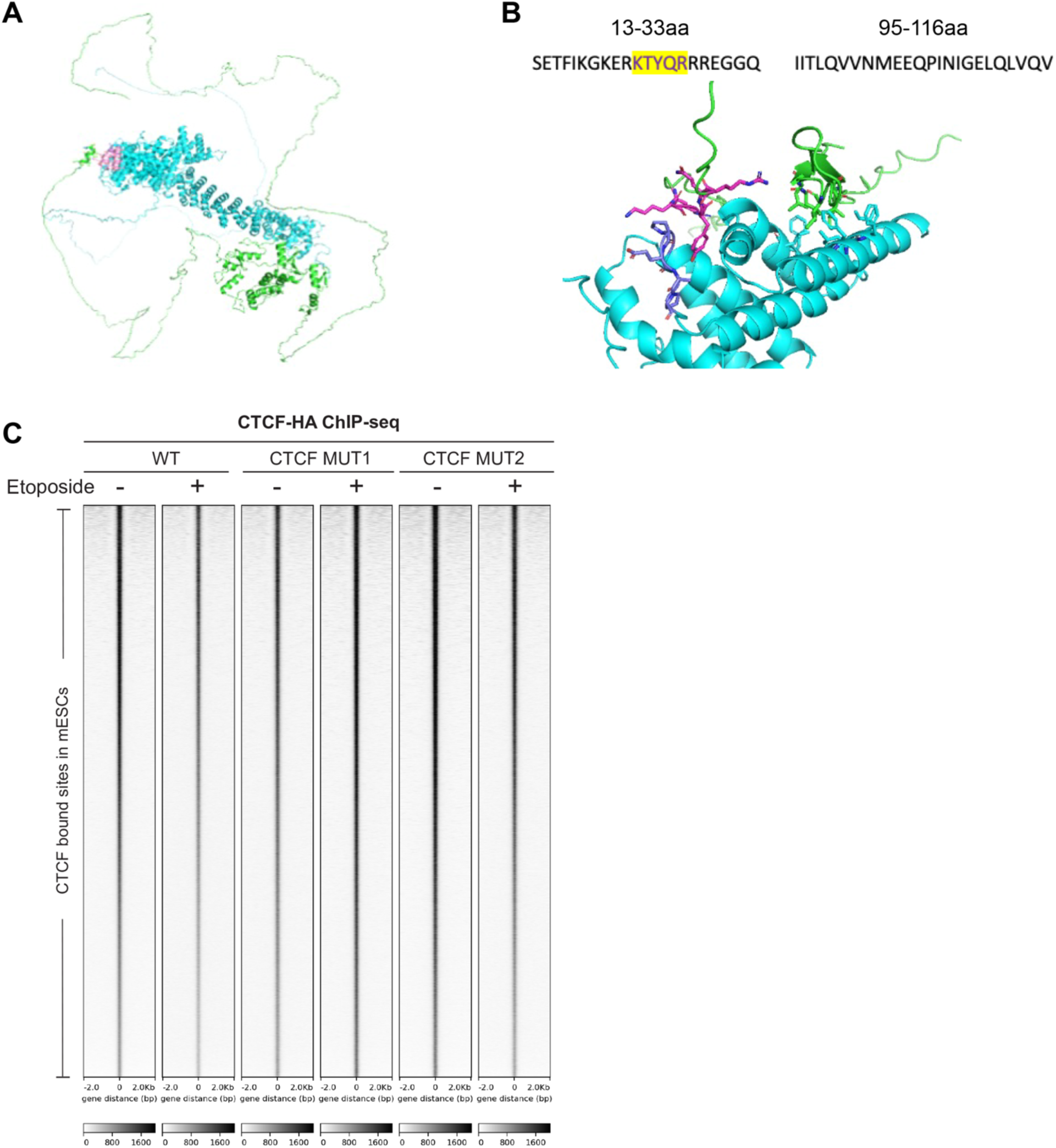
A novel CTCF-PDS5A interacting region on CTCF N-terminus, related to. Fig. 3**. (A)** CTCF and PDS5A interaction prediction model indicating the CTCF N-terminus interacting with PDS5A. The interaction region is highlighted in pink. **(B)** Two relevant interacting regions determined by protein-protein interaction AlphaFold2 software. 13-33 aa (previously reported^17^), and the novel 95-116aa interaction surface were shown. **(C)** Heat maps showing HA-CTCF ChIP-seq read densities at CTCF bound sites in HA-FLAG-CTCF-WT, HA-FLAG-CTCF-MUT1, and HA-FLAG-CTCF-MUT2 mESCs across DMSO versus ETO treatment conditions. ChIP-seq data is from one representative of two biological replicates.

**Fig. S5.**
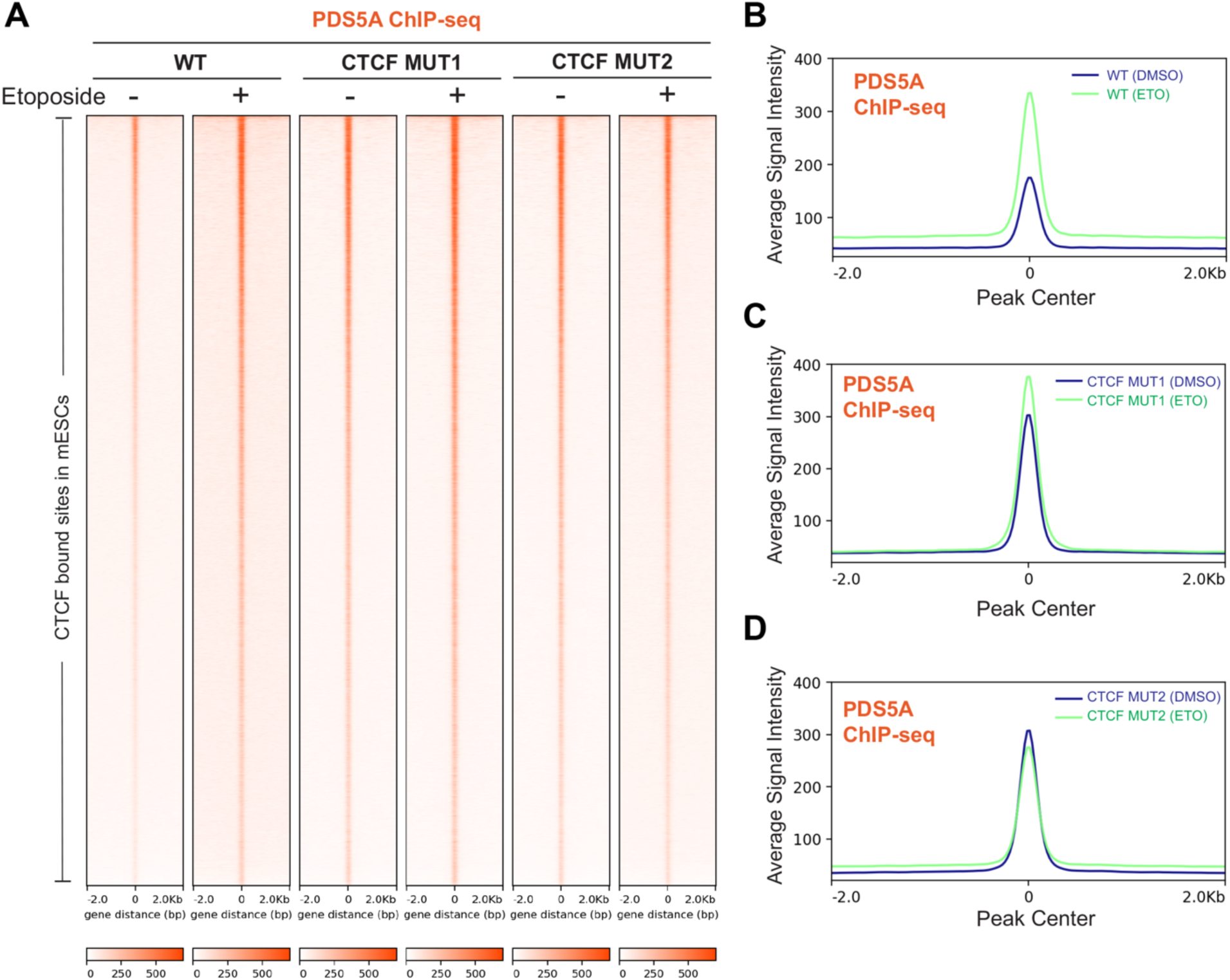
A novel PDS5A-CTCF interaction region in the CTCF N-terminus 95-116 aa is required for PDS5A enrichment at CTCF bound sites, related to. Fig. 3**. (A)** Heat maps showing PDS5A ChIP-seq read densities at CTCF-bound sites in mESCs expressing HA-FLAG-CTCF-WT, HA-FLAG-CTCF-MUT1, and HA-FLAG-CTCF-MUT2 under the conditions of DMSO versus Etoposide treatment. **(B-D)** Average density profile plots showing PDS5A ChIP-seq read densities in CTCF-WT (**B**), CTCF-MUT1 (**C**), and CTCF-MUT2 (**D**) mESCs under DMSO versus Etoposide treatment conditions. ChIP-seq data is from one representative biological replicate across the conditions.

**Fig. S6.**
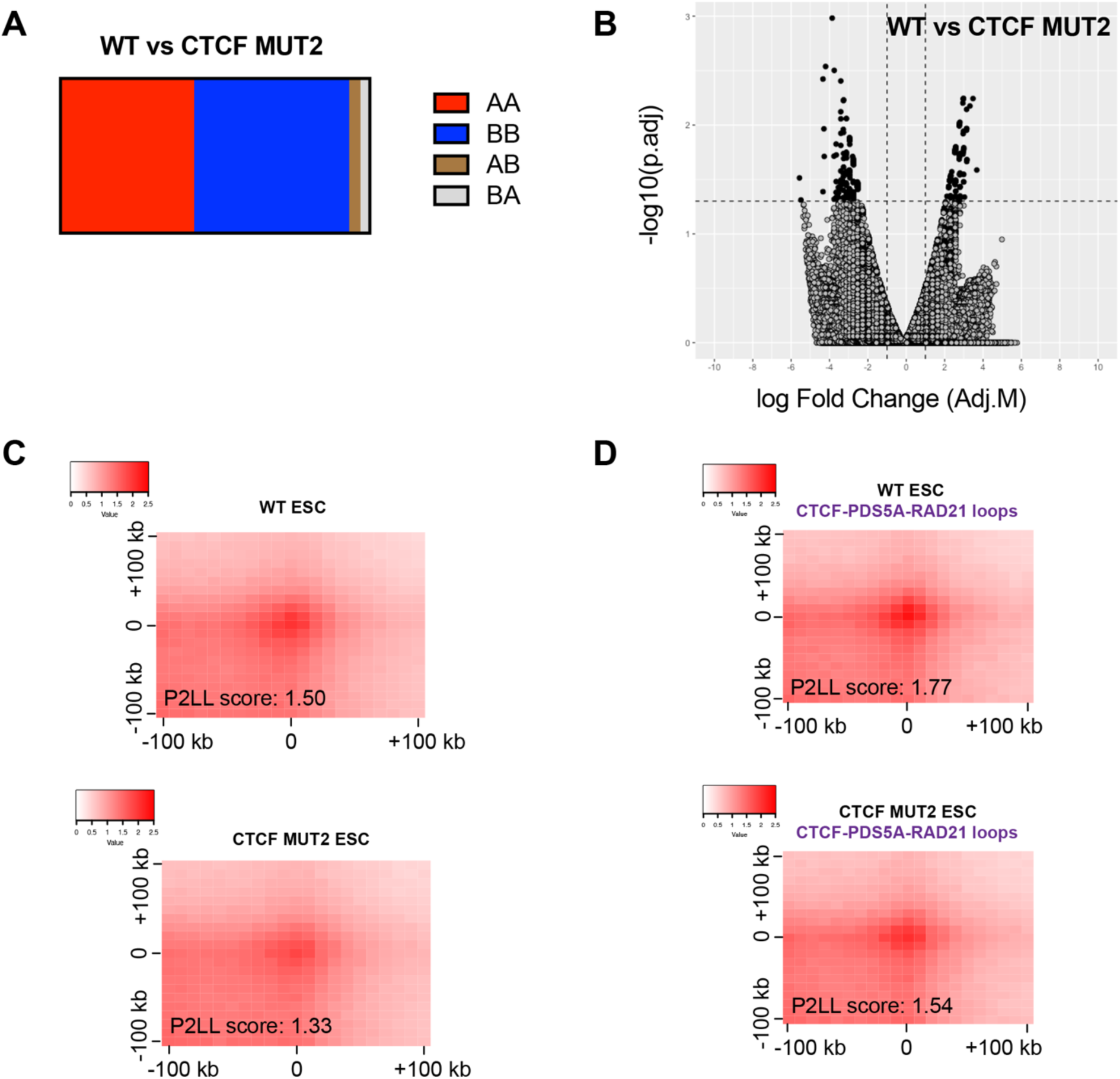
Deletion of 95-116 aa of CTCF N-terminus (CTCF MUT2) impacts genome organization in mESCs, related to. Fig. 4**. (A)** Bar plots showing AB compartments in WT versus CTCF MUT2 mESCs in Micro-C assay (Number of Eigenvector bins - 128 kb were shown). **(B)** Volcano plot of differential loop analysis in WT versus CTCF MUT2 in Micro-C assay. Black circles indicate the differential loops, and gray circles indicate all loops included in the comparative analysis. **(C)** Decrease of loops in Aggregate Peak Analysis (APA) of CTCF MUT2 compared to WT by Micro-C assay in all loops. The resolution of APA is 10 kb. **(D)** Decrease of loops in APA of WT vs CTCF MUT2 mESCs were plotted for the loops with the indicated ChIP-seq signals of CTCF, PDS5A and RAD21 at any region covered by them. The resolution of APA is 10 kb. Micro-C assay represents two biological replicates.

**Fig. S7.**
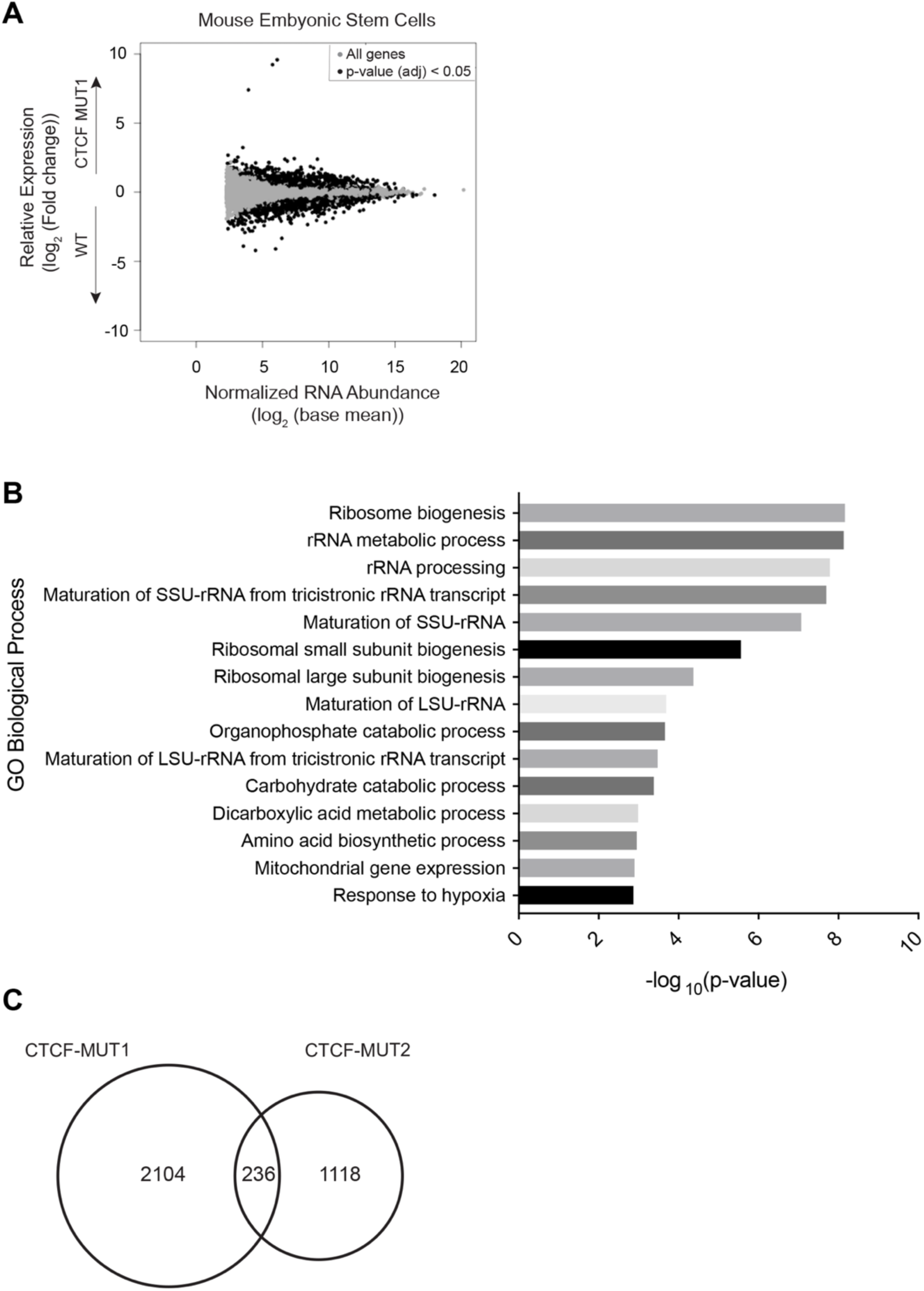
CTCF MUT1 impacts gene expression in mESCs, related to. Fig. 4**. (A)** RNA-seq MA plot of WT vs CTCF MUT1 showing differentially expressed genes in mESCs from four biological replicates. **(B)** Gene Ontology analysis of differentially expressed genes in WT vs CTCF MUT1 RNA-seq in mESCs. **(C)** Venn diagrams showing differentially expressed genes in CTCF-MUT1 versus CTCF-MUT2.

**Fig. S8.**
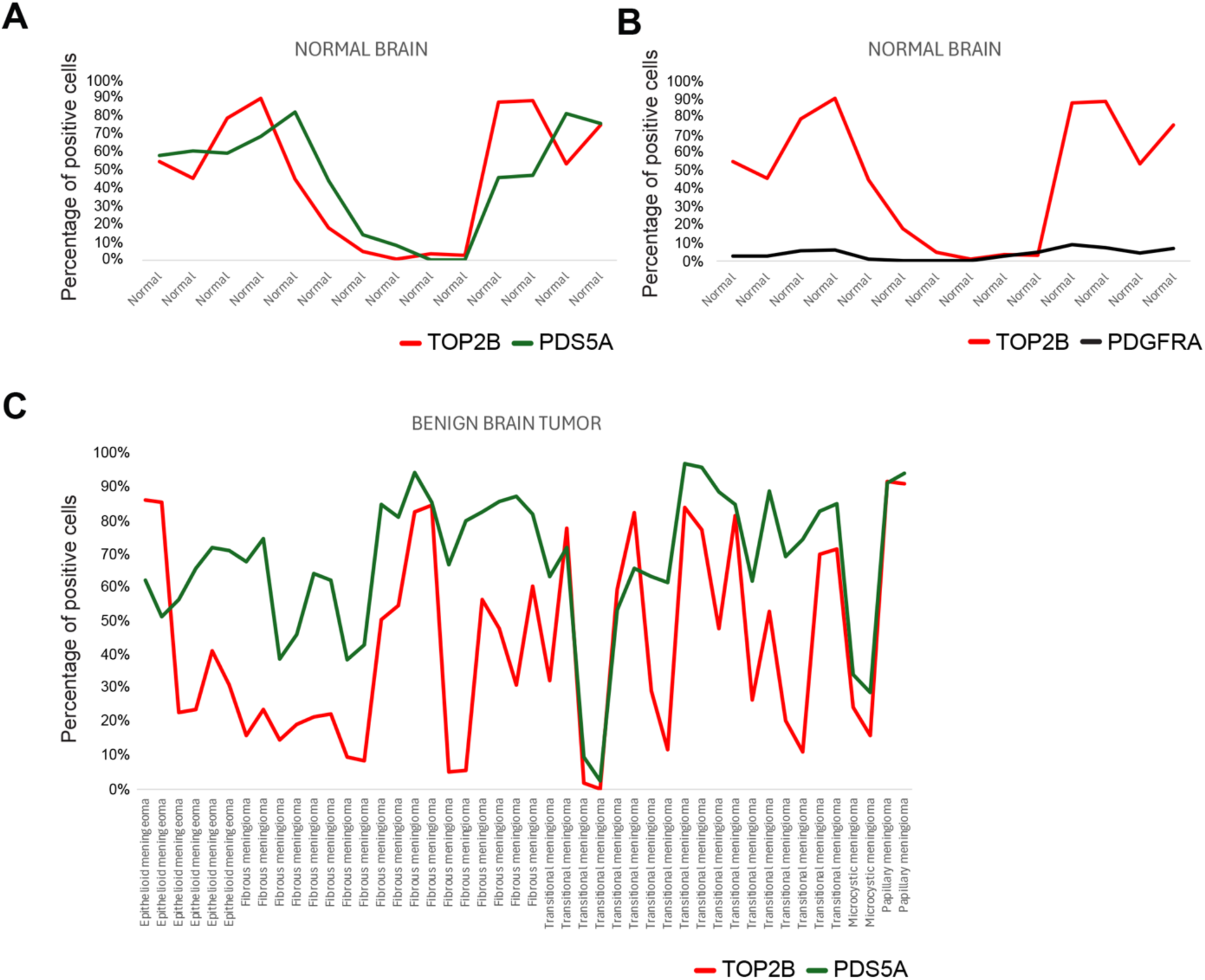
Expressions of PDS5A and TOP2B are variable and correlated in human gliomas, related to. Fig. 5**. (A)** Correlation of TOP2B versus PDS5A expression by Immunohistochemistry in adult normal brain. **(B)** Expression of TOP2B and PDGFRA are not correlated in normal brain, shown as control. **(C)** Correlation of TOP2B versus PDS5A expression by Immunohistochemistry in adult benign brain tumor.

**Fig. S9.**
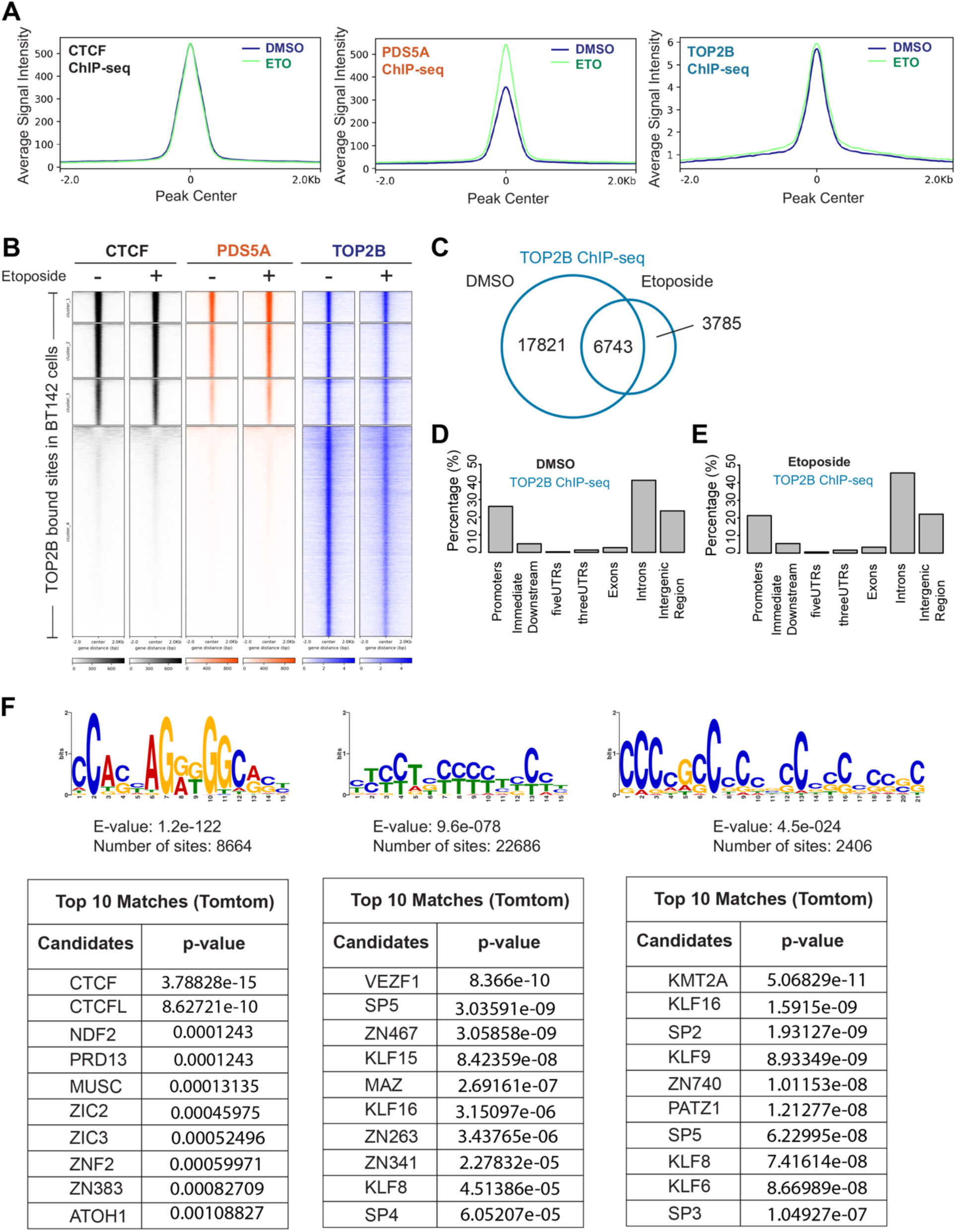
TOP2B activity directly localizes PDS5A at CTCF binding sites in glioma cells, related to. Fig. 6**. (A)** Average density profile plots showing CTCF, PDS5A, and TOP2B ChIP-seq read densities in DMSO versus Etoposide treatment conditions at TOP2B bound sites (shown in Fig. 6E) in BT142 cells. **(B)** Clustered heat maps of CTCF, PDS5A and TOP2B ChIP-seq read densities in DMSO versus ETO treatment conditions in BT142 cells. The clustering was based on default “k-means” clustering that clusters the regions into four groups based on the signal pattern. **(C)** Venn diagram showing TOP2B binding in DMSO versus ETO treatment conditions in BT142 cells. **(D-E)** Genomic distribution of TOP2B binding in BT142 in DMSO (**D**) vs ETO (**E**) treatment conditions. **(F)** Motif analysis of TOP2B binding regions in BT142 cells, along with the table of top matches from Tomtom motif comparison tool indicating the matches to the insulation factors such as CTCF, MAZ, PATZ1, and VEZF1. Motif search in MEME was performed *de novo* until 1000 sites were reached, and the corresponding e-values to the motifs were depicted under the motifs, and the corresponding p-values to the motif matches were depicted in the tables. ChIP-seq data is from one representative of two biological replicates for CTCF and TOP2B, and one replicate for PDS5A. One replicate of TOP2B ChIP-seq has been re-analyzed^45^.

**Fig. S10.**
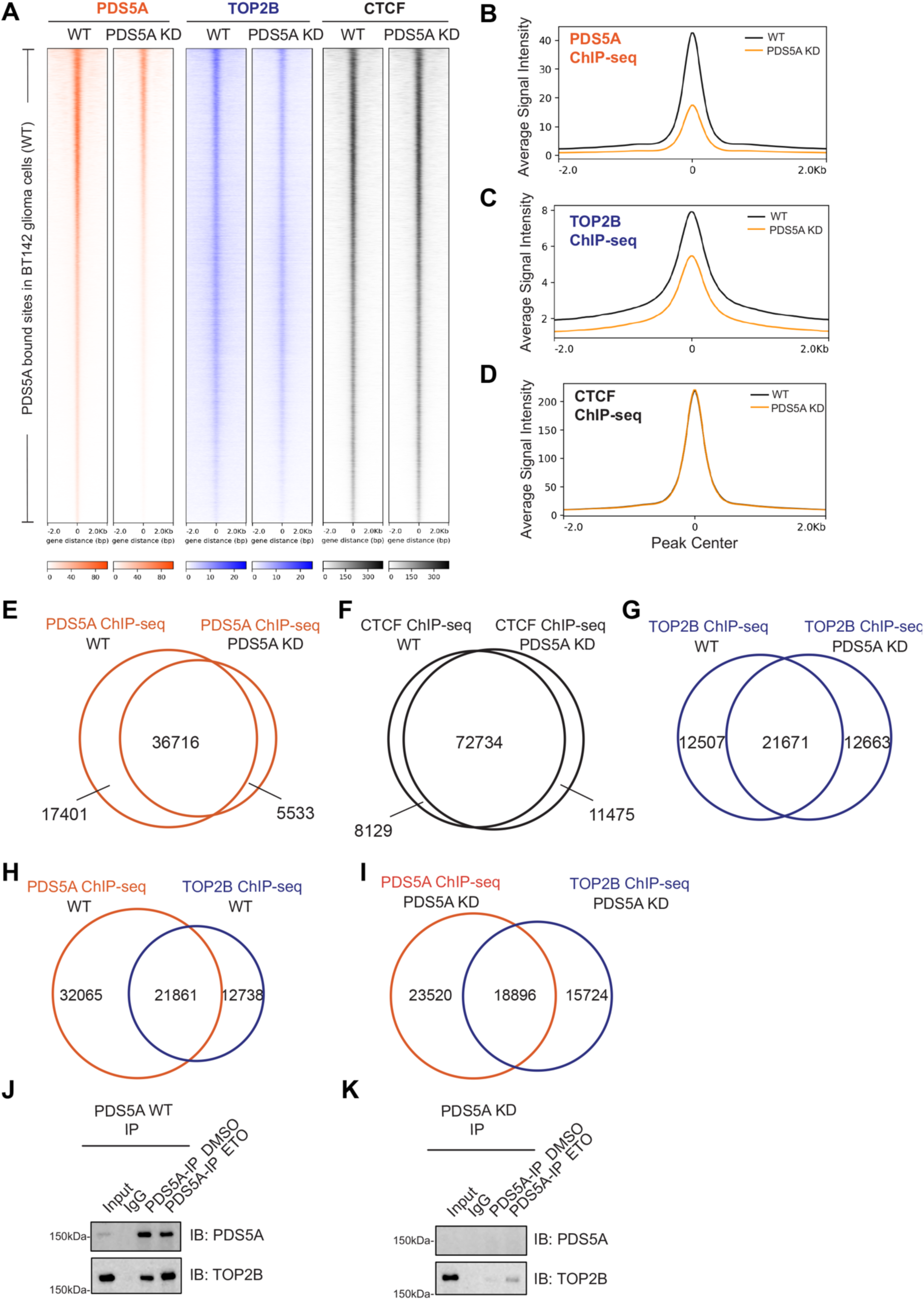
PDS5A knock-down results in reduced occupancy of TOP2B at PDS5A bound sites in BT142 cells, related to. Fig. 7**. (A)** Heat maps showing PDS5A, TOP2B, and CTCF ChIP-seq read densities at PDS5A bound sites in BT142 cells in WT versus PDS5A KD conditions. **(B-D)** Average density profile plots showing PDS5A (**B**), TOP2B (**C**), and CTCF (**D**) ChIP-seq read densities at PDS5A bound sites in BT142 cells in WT versus PDS5A KD conditions. **(E-G)** Venn diagrams indicating the PDS5A (**E**), CTCF (**F**) and TOP2B (**G**) ChIP-seq binding sites in WT versus PDS5A KD conditions in BT142 cells. **(H-I)** Venn diagrams indicating the overlap of PDS5A (orange) and TOP2B ChIP-seq binding sites (blue) in WT condition (**H**), and PDS5A KD condition (**I**) in BT142 cells. ChIP-seq data is from one representative biological replicate for each condition. **(J-K)** TOP2B and PDS5A interaction in BT142 cells under WT (**J**) versus PDS5A KD (**K**) conditions using chromatin fractions isolated from cells treated with DMSO and ETO, respectively, as determined by IP followed by WB (n=2).

**Fig. S11.**
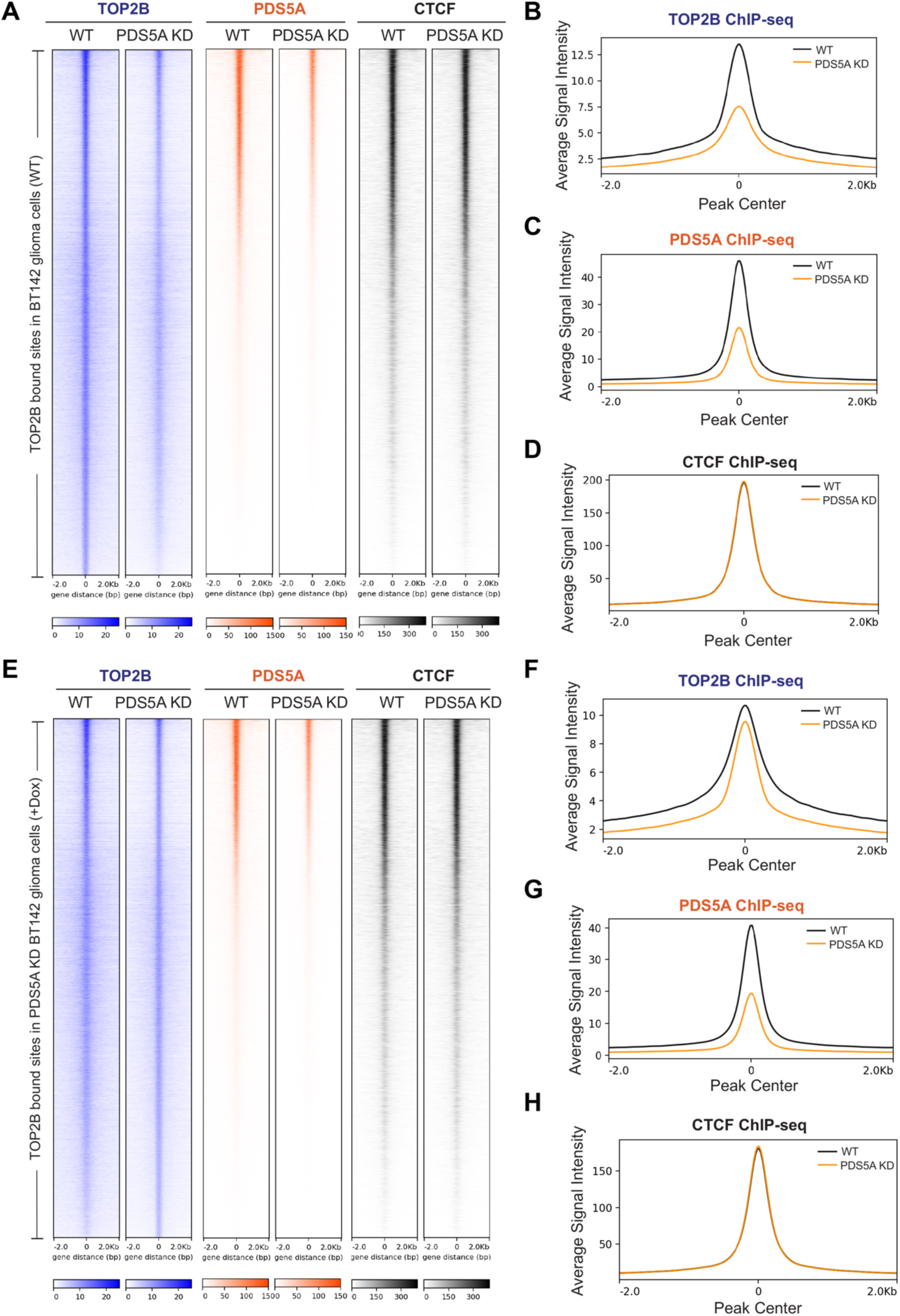
PDS5A knock-down results in the re-localization of TOP2B on chromatin in BT142 glioma cells, related to. Fig. 7**. (A)** Heat maps showing TOP2B, PDS5A and CTCF ChIP-seq read densities at TOP2B bound sites in WT BT142 cells in WT versus PDS5A KD conditions. **(B-D)** Average density profile plots showing TOP2B (**B**), PDS5A (**C**), and CTCF (**D**) ChIP-seq read densities at TOP2B bound sites in WT BT142 cells in WT versus PDS5A KD conditions. **(E)** Heat maps showing TOP2B, PDS5A and CTCF ChIP-seq read densities at TOP2B bound sites in PDS5A KD BT142 cells (+Dox) in WT versus PDS5A KD conditions. **(F-H)** Average density profile plots showing TOP2B (**F**), PDS5A (**G**), and CTCF (**H**) ChIP-seq read densities at TOP2B bound sites in PDS5A KD BT142 cells (+Dox) in WT versus PDS5A KD conditions. ChIP-seq data is from one representative biological replicate for each condition.

**Fig. S12.**
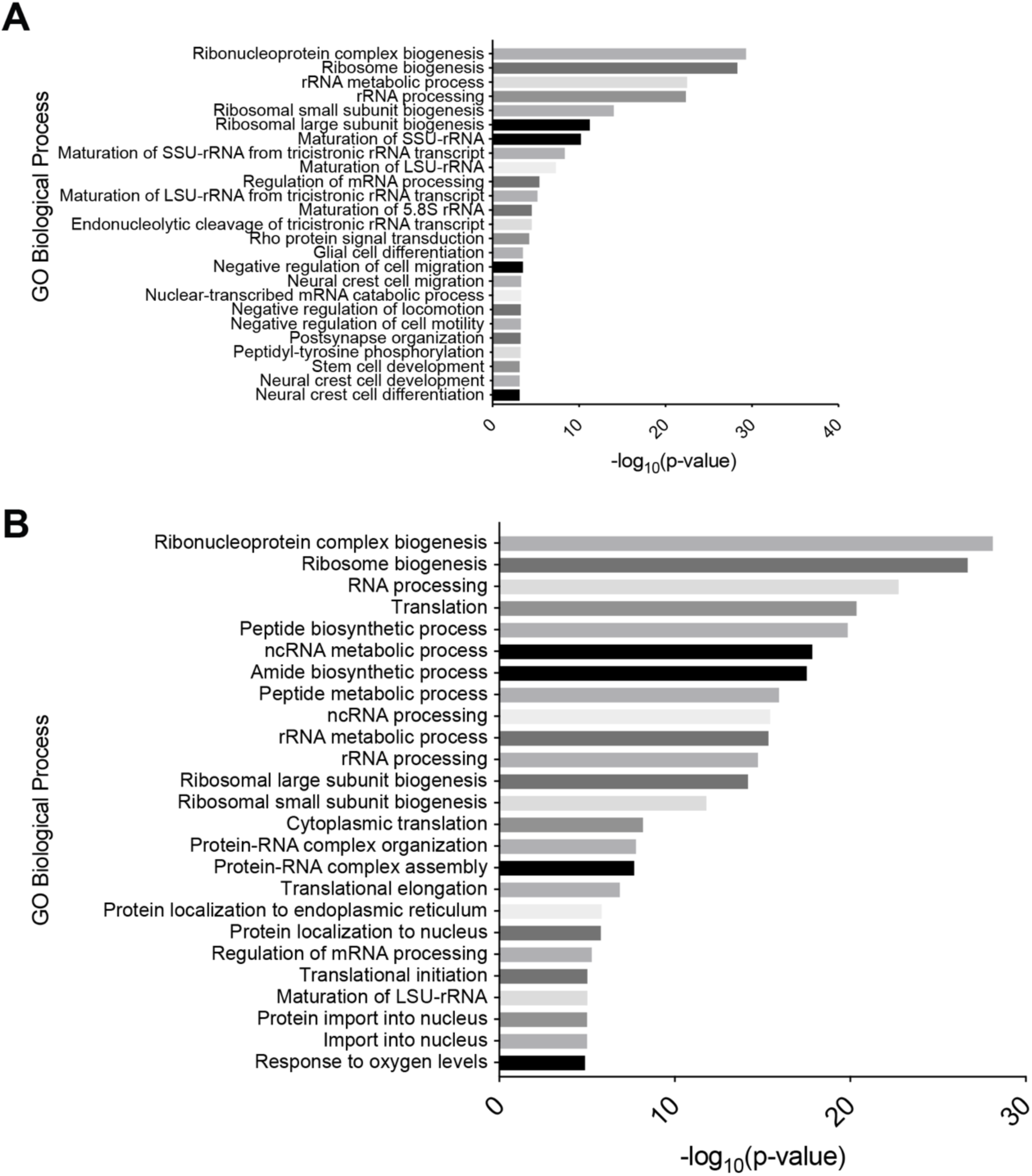
PDS5A knock-down or TOP2B inhibition affects similar biological processes in BT142 glioma cells, related to. Fig. 7**. (A)** Gene ontology analysis of DEGs upon TOP2B inhibition (ICRF-193 treatment) using RNA-seq in BT142 glioma cells. RNA-seq data for control (DMSO) and ICRF-193 treated conditions are from five and seven biological replicates, respectively^45^. **(B)** Gene ontology analysis of DEGs upon PDS5A KD using RNA-seq in BT142 glioma cells from four biological replicates.

---

**Table S1.** Antibodies. Related to Methods (Separate Excel file)

**Table S2.** The Number of Reads in Micro-C Analysis. Related to Fig. 4 and Fig. S6 (Separate Excel file)

**Table S3.** RNA-seq Expression Values in WT vs CTCF MUT1 mESCs. Related to Fig. S7 (Separate Excel file)

**Table S4.** Gene Ontology Analysis in WT vs CTCF MUT1 mESCs. Related to Fig. S7 (Separate Excel file)

**Table S5.** RNA-seq Expression Values in WT vs CTCF MUT2 mESCs. Related to Fig. 4 (Separate Excel file)

**Table S6.** Gene Ontology Analysis in WT vs CTCF MUT2 mESCs. Related to Fig. 4 (Separate Excel file)

**Table S7.** RNA-seq Expression Values in BT142 glioma cells treated with DMSO versus ICRF-193. Related to Fig. 7H (Separate Excel file)

**Table S8.** Gene Ontology Analysis in BT142 glioma cells treated with DMSO versus ICRF-193. Related to fig. S12A (Separate Excel file)

**Table S9.** RNA-seq Expression Values in WT vs PDS5A KD BT142 glioma cells. Related to Fig. 7I (Separate Excel file)

**Table S10.** Gene Ontology Analysis in WT vs PDS5A KD BT142 cells. Related to fig. S12B (Separate Excel file)

**Table S11.** TOP2B IHC quantification. Related to Fig. 5 and Fig. S8 (Separate Excel file)

**Table S12.** PDS5A IHC quantification. Related to Fig. 5 and Fig. S8 (Separate Excel file)

**Table S13.** PDGFRA IHC quantification. Related to Fig. S8 (Separate Excel file)

**Table S14** Brain tumor specifications. Related to Fig. 5 and Fig. S8 (Separate Excel file)

